# Polarized Canine Liver Organoid Monolayers Capture Features of Western Diet-Induced Hepatic Metabolic Reprogramming Across Transcriptomic, Lipidomic, and Biogenic Amine Signatures

**DOI:** 10.64898/2026.11.11.766030

**Authors:** Ryland T. Giebelhaus, Christopher Zdyrski, Paulina de la Mata, Seo Lin Nam, Agnes Bourgois-Mochel, Megan P. Corbett, Haejin Cho, Jessica L. Ward, Emilie Guillot, Eugene Douglass, James J. Harynuk, Jonathan P. Mochel

**Author notes:** <u>Corresponding authors:</u> Ryland T. Giebelhaus, Ph.D., Assistant Professor of Chemistry, Elliott Building Room 313, 3800 Finnerty Road, Victoria, BC V8P 5C2, Canada, Jonathan P. Mochel, DVM, MS, Ph.D., DECVPT, Professor of Systems Pharmacology 501 D.W. Brooks Drive, Athens, GA 30602.

## Abstract

Liver responses to Western-diet (WD) exposure remain incompletely defined, partly because conventional rodent models and immortalized hepatic cell lines incompletely recapitulate species-specific features of liver biology. Here, we established polarized epithelial monolayers from canine liver organoids derived from healthy beagles and beagles exposed to a WD for 8 weeks. We then used this platform to characterize diet-induced hepatic reprogramming and to explore differential responses to two SGLT2 inhibitors, dapagliflozin and bexagliflozin. The monolayers developed reproducible apical-basal polarity, with transepithelial electrical resistance (TEER) values of 123-335 Ω·cm^2^, enabling compartment-resolved sampling. Bulk transcriptomics, lipidomics, and biogenic amine profiling each independently separated WD-derived from healthy monolayers. Periodic acid-Schiff staining provided qualitative support for this diet-associated molecular signature, showing increased carbohydrate-rich material consistent with glycogen accumulation in WD-derived monolayers. By contrast, 24-hour apical exposure to dapagliflozin or bexagliflozin at 20 µM produced no detectable transcriptomic differences and only weak, context-dependent metabolomic separation, which was restricted to the apical compartment in the untargeted metabolomics dataset. These findings establish canine liver organoid-derived polarized monolayers as a tractable *in vitro* platform that captures WD-induced hepatic reprogramming across transcriptomic, lipidomic, and biogenic amine readouts, while providing a foundation for future targeted studies of candidate interventions for diet-associated hepatic dysfunction.

## INTRODUCTION

Western-diet exposure drives hepatic reprogramming across metabolic, inflammatory, and fibrotic pathways (Ishimoto et al., 2013), yet the molecular features of this response remain incompletely captured by conventional experimental models (Cui et al., 2025; Mochel et al., 2024). Clinically, these changes fall within the spectrum of metabolic dysfunction-associated steatotic liver disease (MASLD, formerly NAFLD), now the leading cause of chronic liver disease worldwide (“Erratum,” 2024; European Association for the Study of the Liver (EASL) et al., 2024). There is, therefore, a critical need for establishing tractable preclinical platforms that capture the molecular features of diet-induced hepatic reprogramming and enable their mechanistic interrogation in a translationally relevant setting.

Conventional rodent models and immortalized human hepatocyte lines capture only a limited fraction of species-specific liver biology. Immortalized hepatocyte lines can lose canonical hepatic functions with continued passage, while rodent drug-metabolism and biotransformation pathways differ in important respects from those of humans and large-animal species (Arzumanian et al., 2021; Basit et al., 2022).

Canine liver-derived *in vitro* models provide a useful translational bridge in this context. Dogs develop spontaneous metabolic disorders, including obesity-associated hepatic steatosis, insulin resistance, and dyslipidemia, under shared environmental and dietary exposures that more closely approximate human cardiometabolic disease than many experimentally induced rodent models (Mochel et al., 2024; Tvarijonaviciute et al., 2012; Yin et al., 2012). Three-dimensional organoids derived from primary adult tissue have emerged over the past decade as tractable *in vitro* platforms that preserve donor cellular identity, develop apical-basal polarity, and recapitulate key functional responses of the parent epithelium (Gabriel, Zdyrski, et al., 2024). Liver organoids have now been derived from multiple species, but canine liver organoid systems remain comparatively rare and have been applied largely to inherited copper-storage disorders, most prominently the COMMD1-deficient phenotype (Wilson’s disease), rather than to acquired, diet-induced hepatic disease (Nantasanti et al., 2015). A persistent challenge for liver-derived organoid systems is the limited differentiation of organoid epithelium toward a mature hepatocyte phenotype. Because canine liver organoids retain a prominent cholangiocyte-like signature under standard expansion conditions, downstream functional readouts must be interpreted in light of the relative contributions of hepatocyte-like and cholangiocyte-like populations (Gabriel, Lincoln, et al., 2024). When grown on Transwell membranes, organoid-derived epithelia form polarized monolayers amenable to compartment-resolved sampling, allowing apical drug exposure and separate assessment of apical and basal metabolic responses (Gabriel et al., 2022). This format is well suited to multiplatform metabolomic profiling, in which untargeted metabolomics, lipidomics, and biogenic amine profiling can provide a more comprehensive molecular characterization than any single analytical platform alone.

Whether organoids derived from donor tissue exposed to defined dietary conditions preserve exposure-associated phenotypes after *in vitro* expansion remains insufficiently characterized, particularly for Western-diet-induced hepatic reprogramming. In this context, the *in vivo* phenotype reflects sustained metabolic reprogramming that may be attenuated, lost, or reprogrammed during tissue dissociation, serial passage, cryopreservation, and expansion on Transwell inserts. Demonstrating phenotype retention therefore requires convergent evidence across morphological, transcriptomic, and metabolic readouts. To our knowledge, no prior study has addressed this question in canine liver organoids using an integrated multi-omics design.

Sodium-glucose cotransporter 2 (SGLT2) inhibitors provide a relevant pharmacologic context for this question. Originally developed to improve glycemic control in type 2 diabetes mellitus by blocking glucose reabsorption in the proximal renal tubule, the gliflozin class has since demonstrated cardiovascular, renal, and weight-related benefits in large outcome trials (Birkeland et al., 2019; Butler et al., 2021; Inzucchi et al., 2020; Kosiborod et al., 2023; McMurray et al., 2019; Neal et al., 2017; Packer et al., 2020; Persson et al., 2018). A growing literature also supports broader effects on lipid metabolism, systemic inflammation, and energy balance (Packer et al., 2020). Hepatic responses are relevant because MASLD is prevalent in the same cardiometabolic populations for whom SGLT2 inhibitors are commonly prescribed. However, because SGLT2/SLC5A2 is expressed predominantly in the renal proximal tubule rather than being a canonical hepatocyte target, hepatic effects of SGLT2 inhibitors are generally expected to arise indirectly through renal and systemic metabolic adaptations, or potentially through noncanonical activity at related transporters or enzymes. Whether the second-generation gliflozin, bexagliflozin, produces a hepatic metabolic fingerprint distinguishable from that of dapagliflozin has not been formally addressed in any species.

The present study had two related objectives. First, and primarily, we asked whether polarized monolayers derived from canine liver organoids from two healthy beagles and two beagles fed a Western diet for 8 weeks recapitulate diet-induced hepatic reprogramming across complementary histologic, transcriptomic, and metabolomic readouts. Second, as a purely exploratory objective, we used the same platform to assess whether 24-hour apical exposure to dapagliflozin or bexagliflozin produced metabolomic changes distinguishable from vehicle control. The drug comparison was designed as a hypothesis-generating component rather than a confirmatory efficacy assessment, given the limited number of donor-derived organoid lines per dietary group and the supratherapeutic 20 µM exposure, which is approximately two orders of magnitude above clinically achieved unbound plasma concentrations for either compound.

## MATERIALS AND METHODS

### Organoid generation

Liver tissue samples from healthy dogs were obtained post-mortem from two necropsy cases, in compliance with Standard Operating Procedures (SOPs) approved by the Institutional Animal Care and Use Committee at Marshall BioResources. All procedures adhered to USDA Animal Welfare Regulations. The two healthy liver organoid lines have previously been described from our group (Gabriel, Lincoln, et al., 2024). The two samples representing the Western diet (WD) group were collected laparoscopically from research beagles that had been maintained on a WD for 8 weeks, at Iowa State University (IACUC-19-337), as previously described by our consortium (Mason et al., 2025). Donor demographics are summarized in **Table 1**.

**Table 1:**
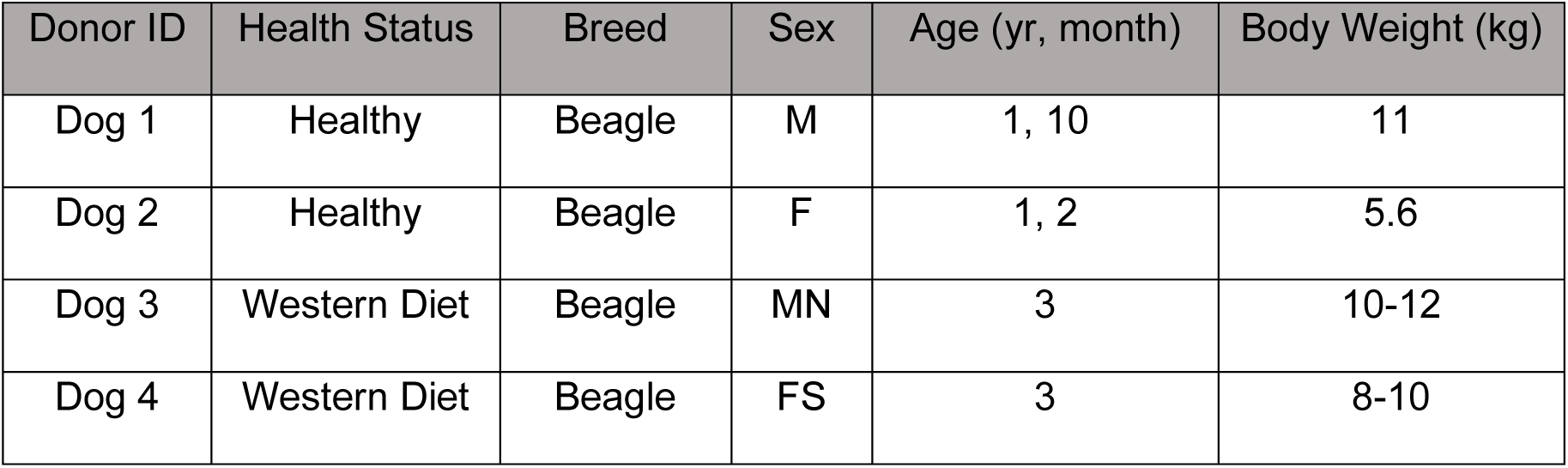
Patient details including breed, sex, age, and body weight. Female spayed = FS, Male neutered = MN. All details regarding the composition of the WD can be found in Mason et al. (2025).

Adult stem cell-derived organoids were generated and expanded from these tissues using previously described methods (Gabriel et al., 2022). In brief, tissues were subjected to CCS digestions, resuspended in Matrigel, and plated with remaining liver tissue. The organoid media (CMGF+) was prepared as previously described (Zdyrski et al., 2025) and added to the wells after the Matrigel had polymerized in the incubator. Samples were passaged to expand the cultures, cryopreserved, and subsequently thawed for experimental use.

### Transwell seeding

Thawed organoids were expanded and then dissociated to single cells and seeded onto 24-well Transwell inserts (0.33 cm^2^ growth area; Corning), as previously described (Gabriel et al., 2022). Transepithelial electrical resistance (TEER) was measured every 1-2 days using a Millicell ERS-2 Epithelial Volt-Ohm Meter (Millipore; MERS00002), with daily calibration to ∼1,000 Ω against the Test Electrode (MERSSTX04) and measurements taken with chopstick probes (MERSSTX01). On each plate, one Transwell coated with Matrigel but receiving no cells served as a blank; sample resistance was corrected by subtracting the blank value before area correction (TEER [Ω·cm²] = [Raw – Blank] × 0.33). Once TEER plateaued, expansion medium (CMGF+ R/G) was exchanged on both apical and basal compartments for differentiation medium (CMGF+).

### Drug exposure

Dapagliflozin propanediol monohydrate (Selleckchem; S5566) and bexagliflozin (MedChemExpress; HY-17604/CS-5702) were dissolved in DMSO at 20 mM and diluted in CMGF+ medium to a final working concentration of 20 µM. We acknowledge that the 20 µM screening concentration is markedly supratherapeutic relative to clinical exposure. At therapeutic doses, both agents reach low sub-micromolar total plasma Cmax values: approximately 62 ng/mL (∼150 nM) for dapagliflozin at 5 mg (De Bruin et al., 2016), with approximately dose-proportional exposure across the therapeutic range (Kasichayanula et al., 2014), and approximately 134 ng/mL (∼290 nM) for bexagliflozin at 20 mg (TheracosBio, LLC, 2025). Given plasma protein binding of approximately 91% for dapagliflozin (AstraZeneca Pharmaceuticals LP, 2024)) and approximately 93% for bexagliflozin (TheracosBio, LLC, 2025), 20 µM exceeds the total plasma Cmax of either drug by approximately two orders of magnitude, and the estimated unbound Cmax by approximately three. This supraphysiological exposure was used deliberately to maximize the probability of detecting compound-specific metabolic perturbations within the limited dynamic range of an *in vitro* screen, with the explicit caveat that any signal identified here cannot be directly extrapolated to clinical exposure and would require a dose-response in a subsequent study to be interpreted pharmacologically.

Compounds were incubated on the apical side of the Transwell for a total of 24 hours at 37 °C, 5% CO_2_. Each donor contributed three technical replicates per treatment arm (vehicle, dapagliflozin, bexagliflozin); biological replication was therefore n = 2 donors per health state and per arm, and the technical replicates do not provide independent biological power. At the end of the incubation, apical (200 µL) and basal (700 µL) supernatants were collected separately and stored at -80°C until shipment for metabolomics analysis.

### Fixation and staining

For hematoxylin and eosin (H&E) staining, organoid medium was removed, and 500 µL of Formalin-acetic acid-alcohol (FAA, composition in Gabriel et al. 2022) made from Ethanol (VWR: 71002-512), Acetic Acid (Fisher; A38500), Formaldehyde (Fisher; F79P4), and MilliQ water, was added to each well (Gabriel et al., 2022). After 24 hours, FAA was replaced with 70% ethanol, which was made from diluting 100% ethanol, and samples were paraffin-embedded and stained for H&E at Iowa State University. Tissues were fixed in formalin and paraffin-embedded according to standard histology procedures. Slides were cut and periodic-acid Schiff (PAS) stained with and without diastase digestion at the University of Georgia Histopathology core and were then imaged using an Olympus BX41 Microscope with a BioVid 4k HD camera (LW Scientific). Images were color-corrected, and scale bars were added with ToupView, version x64 131 4.11.23945.20231121 (LW Scientific).

### Transmission electron microscopy

After Transwell monolayers were established, media was removed from the apical and basal compartments. Transwell inserts were then filled with 1% paraformaldehyde and 3% glutaraldehyde in phosphate-buffered saline (PBS) and stored at 4°C prior to processing. The samples were fixed for 48 hours at 4°C in 1% paraformaldehyde, 3% glutaraldehyde in 0.1 M sodium cacodylate buffer, pH 7.2. After washing in a cacodylate buffer three times for 10 min each, samples were post-fixed with 1% osmium tetroxide in 0.1M sodium cacodylate buffer at room temperature for 1 hour. Next, the samples were washed with deionized water three times for 15 minutes each, and then bloc stained using 2% uranyl acetate in distilled water for 1 hour. Samples were washed for 10 minutes in distilled water, dehydrated for 1 hour in each step of a graded ethanol series (25, 50, 70, 85, 95, 100%), dehydrated further with three changes of pure acetone for 15 minutes each, and then infiltrated with EmBed 812 formula (hard) for EPON epoxy resin (Electron Microscopy Sciences, Hatfield, PA) with graded ratios of resin to acetone until fully infiltrated with pure epoxy resin (3:1, 1:1, 1:3, pure) for 6-12 hours per step. The samples were then placed into BEEM capsules and polymerized at 70°C for 48 hours. Using a Leica UC6 ultramicrotome (Leica Microsystems, Buffalo Grove, IL), 1.5 µm thick sections were made and stained with EMS Epoxy stain (a blend of toluidine blue-O and basic fuchsin). Thin sections were made at 50 nm and collected onto single slot carbon film grids. Finally, TEM images were captured using a 200 kV JEOL JSM 2100 scanning transmission electron microscope (Japan Electron Optics Laboratories, Peabody, MA) with a GATAN One View 4K camera (Gatan inc., Pleasanton, CA).

### RNA extractions

Organoids were preserved for bulk RNA-sequencing by first harvesting the apical and basal supernatants from the Transwell. Then, 200 µL of Trizol (Invitrogen; 15596026) was added to the apical side of each Transwell and incubated for 15 minutes at room temperature. After incubation, the apical sides were pipette mixed, pooled together, and a final rinse of a fresh 200 µL of Trizol was used per condition. Samples were then stored at -80°C. In general, RNA extractions were carried out as previously described (Nicholson et al., 2026). The organoids samples were thawed and more Trizol was added to reach 1 mL total per tube, and each sample was briefly vortexed. Organoid samples were left at room temperature for five minutes before centrifuging at 12,000 x g (4°C) for 10 minutes. The supernatant was transferred to a microcentrifuge tube and 200 µL of chloroform (Alfa Aesar; J67241) was added to the samples. The samples were vigorously mixed by shaking for 20 seconds. After sitting at room temperature for 2-3 minutes, samples were spun at 10,000 x *g* for 18 minutes (4°C). The aqueous top layer was collected and moved to a sterile RNase-free tube before adding an equal amount of 100% RNA-free ethanol. Up to 700 µL were loaded in a Qiagen RNeasy column and collection tube (RNeasy Mini kit; Qiagen; 74104). Samples were centrifuged at 8,000 x *g* for 30 seconds, the waste in the collection tube was discarded, and DNase treatment was performed according to Qiagen’s protocol (Qiagen; 79254). In a new collection tube, 500 µL buffer RPE was added to the column, and the samples were spun at the same settings as before. After discarding the flow-through, this step was repeated, but samples were now spun for 2 minutes at 8,000 x *g*. The flow-through was discarded again, and the samples were centrifuged at the same settings for 1 minute. 50 µL of RNase-free water (Sigma; W4502-50ML) was added to the column and incubated for 2 minutes. Samples were centrifuged twice at 8,000 x *g* for 1 minute. RNA concentration was determined with a Nanodrop ND-1000 Spectrophotometer (Thermo Fisher Scientific), and samples were kept at -80°C until shipped.

### Strand-specific RNA library preparation and sequencing

RNA was quantified on a Qubit 2.0 Fluorometer (Life Technologies, Carlsbad, CA, USA) and RNA integrity assessed on an Agilent TapeStation 4200 (Agilent Technologies, Palo Alto, CA, USA). Strand-specific RNA sequencing libraries were prepared using the NEBNext Ultra II Directional RNA Library Prep Kit for Illumina (NEB) following the manufacturer’s instructions (NEB, Ipswich, MA, USA). Briefly, the enriched RNAs were fragmented for 8 minutes at 94°C. First strand and second strand cDNA were subsequently synthesized. The second strand of cDNA was marked by incorporating dUTP during the synthesis. cDNA fragments were adenylated at 3’ends, and indexed adapter was ligated to cDNA fragments. Limited cycle PCR was used for library enrichment. The incorporated dUTP in second strand cDNA quenched the amplification of second strand, which helped to preserve the strand specificity. The sequencing library was validated on the Agilent TapeStation (Agilent Technologies, Palo Alto, CA, USA) and quantified by using Qubit 2.0 Fluorometer (ThermoFisher Scientific, Waltham, MA, USA) as well as by quantitative PCR (KAPA Biosystems, Wilmington, MA, USA). The sequencing libraries were multiplexed and clustered onto a flowcell on the Illumina NovaSeq instrument according to manufacturer’s instructions. The samples were sequenced using a 2×150bp Paired End (PE) configuration. Image analysis and base calling were conducted by the NovaSeq Control Software (NCS). Raw sequence data (.bcl files) generated from Illumina NovaSeq was converted into fastq files and de-multiplexed using Illumina bcl2fastq 2.20 software. One mismatch was allowed for index sequence identification.

### RNA analysis

Raw sequencing reads were assessed for quality using FastQC and FASTP (Chen et al., 2018). High-quality reads were pseudoaligned to the canine reference transcriptome (CanFam3.1 cDNA) using Kallisto (Bray et al., 2016). Transcript-level abundances were quantified and aggregated to the gene level using tximport (Soneson et al., 2015), and Ensembl gene IDs were converted to gene symbols using biomaRt (Durinck et al., 2005, 2009). Gene-level expression values were transformed to log2(TPM+1) and used for principal component analysis (PCA). Differential expression analysis was conducted using DESeq2 (Love et al., 2014), comparing WD and healthy control samples. Genes with a total count < 10 across all samples were filtered out and genes with an adjusted *p*-value < 0.05 were considered differentially expressed. Heatmaps were generated from log2-transformed TPM values using the heatmap.2 function in the gplots R package (v 4.4.0).

### Metabolomics analyses

All metabolomic analyses were performed by the West Coast Metabolomics Center at the University of California, Davis (Davis, CA, USA), on aliquots shipped frozen from our laboratory.

#### General metabolism

Samples were extracted using the extraction procedure by Matyash et al. (2008), which includes MTBE, MeOH, and H_2_O (Matyash et al., 2008). The organic (upper) phase was dried down and submitted for resuspension and injection onto the LC, while the aqueous (bottom) phase was dried down and submitted for derivatization for GC. Samples were shaken at 30°C for 1.5 hours. Then, 91 μL of MSTFA + FAMEs were added to each sample, and tubes were shaken at 37°C for 0.5 hours to complete the derivatization. Samples were then vialed, capped, and injected onto the instrument. A 7890A GC coupled with a LECO time of flight mass spectrometer (TOFMS) was used for the procedure. Then, 0.5 μL of the derivatized sample was injected using a splitless method onto a RESTEK RTX-5SIL MS column (30 m × 0.25 mm inner diameter with 0.25 μm film thickness) with an Integra-Guard at 275°C with a helium flow of 1 mL/min. The GC oven was set to hold at 50°C for 1 minute, then ramped up to 20°C/min to 330°C and held for 5 minutes. The transfer line was set to 280°C, while the EI ion source was set to 250°C. The mass spectrometry parameters collected data from 85 m/z to 500 m/z at an acquisition rate of 17 spectra/second.

All compounds detected were tentatively identified to the Metabolomics Standards Initiative (MSI) Level 2 with a spectral library match score of 800 or higher and retention indices within ± 2,000 units (Fiehn retention index), corresponding to about ± 2.0 s in retention time (Sumner et al., 2007).

#### Lipidomics

Samples were extracted using the extraction procedure by Matyash et al. (2008), which includes MTBE, MeOH, and H_2_O (Matyash et al., 2008). The organic (upper) phase was dried down and resuspended for injection onto the LC, while the aqueous (bottom) phase was dried down and submitted for derivatization for GC. The samples were then resuspended with 110 μL of a solution of 9:1 methanol:toluene and 50 ng/mL CUDA. Samples were then shaken for 20 seconds, sonicated for 5 minutes at room temperature, and centrifuged for 2 minutes at 16,100 rcf.

Thirty-three μL of samples were aliquoted into a vial with a 50 μL glass insert for positive and negative mode lipidomics. The samples were then loaded onto an Agilent 1290 Infinity LC stack. Aliquots were injected in both positive ionization and negative ionization modes. The positive mode was run on an Agilent 6546 with a scan range of m/z 120-1200 Da, an acquisition speed of 2 spectra/s, and a mass resolution of 10,000. Positive mode had between 0.5 and 2 μL injected onto an Acquity Premier BEH C18 1.7 μm, 2.1 x 50 mm column. In positive ionization mode, mobile phase A consisted of 60:40 v/v acetonitrile:water with 10 mM ammonium formate and 0.1% formic acid, and mobile phase B consisted of 90:10 v/v isopropanol:acetonitrile with 10 mM ammonium formate with 0.1% formic acid. The gradient used was 0 min 15% (B), 0.75 min 30% (B), 0.98 min 48% (B), 4.00 min 82% (B), 4.13-4.50 min 99% (B), 4.58-5.50 min 15% (B) with a flow rate of 0.8 mL/min.

Another aliquot was run in negative mode on an Agilent 1290 Infinity LC stack and injected onto the same column, with the same gradient as in positive mode, using an Agilent 6550 QTOF mass spectrometer. In negative ionization mode, mobile phase A was 60:40 v/v acetonitrile:water with 10 mM ammonium acetate, and mobile phase B was 90:10 v/v isopropanol:acetonitrile with 10 mM ammonium acetate. The acquisition rate was two spectra per second with a scan range of m/z 60-1200 Da, and a mass resolution of 20,000. All annotated lipids were putatively identified to MSI Level 2 against the LipidBlast spectral library (Kind et al., 2013). Manual curation and inspection of all spectra was performed to increase certainty in identification.

#### Biogenic amines

Sample extraction for biogenic amines consisted of a liquid-liquid extraction method with MTBE, methanol, and water, creating a biphasic partition (Matyash et al., 2008). The polar phase was then dried down to completion and run on a Waters Premier Acquity BEH Amide column. A short 4-minute liquid chromatography method was used for the separation of polar metabolites from a starting condition of 100% LC-MS H_2_O with 10 mM ammonium formate and 0.125% formic acid to an end condition of 100% ACN:H_2_O 95:5 (v/v) with 10 mM ammonium formate and 0.125% formic acid. A Sciex Triple-ToF scanned from 50-1500 m/z with MS/MS collection from 40-1000, selecting the top five ions per cycle. Data processing was done with MS-Dial using an MZ-RT list for annotations, in addition to a library for MS/MS matching. All annotated biogenic amines were putatively identified to MSI Level 2 using both in-house MS/MS databases and the NIST database, followed by manual curation and inspection of all mass spectra.

#### Quality Control

Sample injection order was randomized across all platforms. Pooled QC samples were injected every 11 injections for all analyses. QC samples clustered tightly in PCA with no drift or obvious batch effects in all cases (Evans et al., 2020; Giebelhaus et al., 2026). For general metabolism data all metabolites in each sample were normalized to the total abundance of all annotated metabolites (MSI Level 2) to account for instrument drift. For lipidomics and biogenic amines, all metabolites were normalized to the total abundance of internal standards in that sample.

#### Statistics

For metabolomic analyses, normalized peak tables were uploaded into MATLAB® (R2023b, The Mathworks Inc., Natick, MA) and labeled by donor, health state, compartment, and treatment. PLS_Toolbox v9.0 (Eigenvector Research, Manson, WA, USA) was used for autoscaling and principal component analysis (PCA). PCA was used as an unsupervised, exploratory visualisation step throughout; no supervised classification was attempted.

PERMANOVA was performed on the peak tables containing all variables for hypothesis generation and exploratory analysis to identify PCA models where there was considerable separation between groups. Given the small sample size (N = 4 with 3 technical replicates), PERMANOVA was applied for hypothesis generation and exploratory analysis rather than a confirmatory statistical test. PERMANOVA on autoscaled peak tables was performed in R v4.4.1 with the vegan package v2.6-4, using Euclidean distance and 99,999 permutations and an α threshold of 0.05. Comparisons which were significant by PERMANOVA were subjected to Permutational Analysis of Multivariate Dispersions (PERMDISP) to compare within-group dispersion (Anderson, 2006). This was performed with the vegan package v2.6-4 in R (v4.4.1), using spatial medians calculated from the same Euclidean distance matrix used for PERMANOVA, with significance evaluated using 99,999 permutations.

Variables driving between-group separation were identified by Cluster Resolution Feature Selection (FS-CR) (Adutwum et al., 2017; Armstrong et al., 2021; Sinkov & Harynuk, 2011). FS-CR sequential backward elimination and forward selection was applied across 100 stratified permutations of each dataset, with only features retained in more than 85 % of permutations carried forward, controlling for overfitting. The cluster-resolution metric, an inter-versus intra-cluster distance ratio in PCA space, was used to score class separation.

For a focused pathway-level analysis on the general metabolomics platform, eleven *a priori* metabolites of interest spanning the TCA cycle, glycolysis, amino-acid biosynthesis, and cholesterol/bile-acid metabolism were extracted from the annotated feature table and compared between treatment arms within each health state. Each metabolite was tested by two-tailed unpaired *t*-test, with Benjamini-Hochberg FDR correction applied across the full untargeted matrix of 756 variables. Fisher ratios were computed alongside the *t-*statistics.

## RESULTS

### Organoid growth

Four canine liver organoid lines — two from healthy adult beagles (one F, one M) and two from beagles maintained on a WD for 8 weeks (one MN, one FS) — were established as 3D cultures and successfully transitioned to polarized Transwell monolayers for all donors (**Figure 1 a-c**). On the final day of the experiment, area-corrected TEER values ranged from 123-335 Ω·cm² across donors (**Figure 1d**). These values are at the low end of those reported for tight hepatic and biliary epithelia and are consistent with a cholangiocyte-enriched monolayer phenotype, which is addressed further in the Discussion.

**Figure 1:**
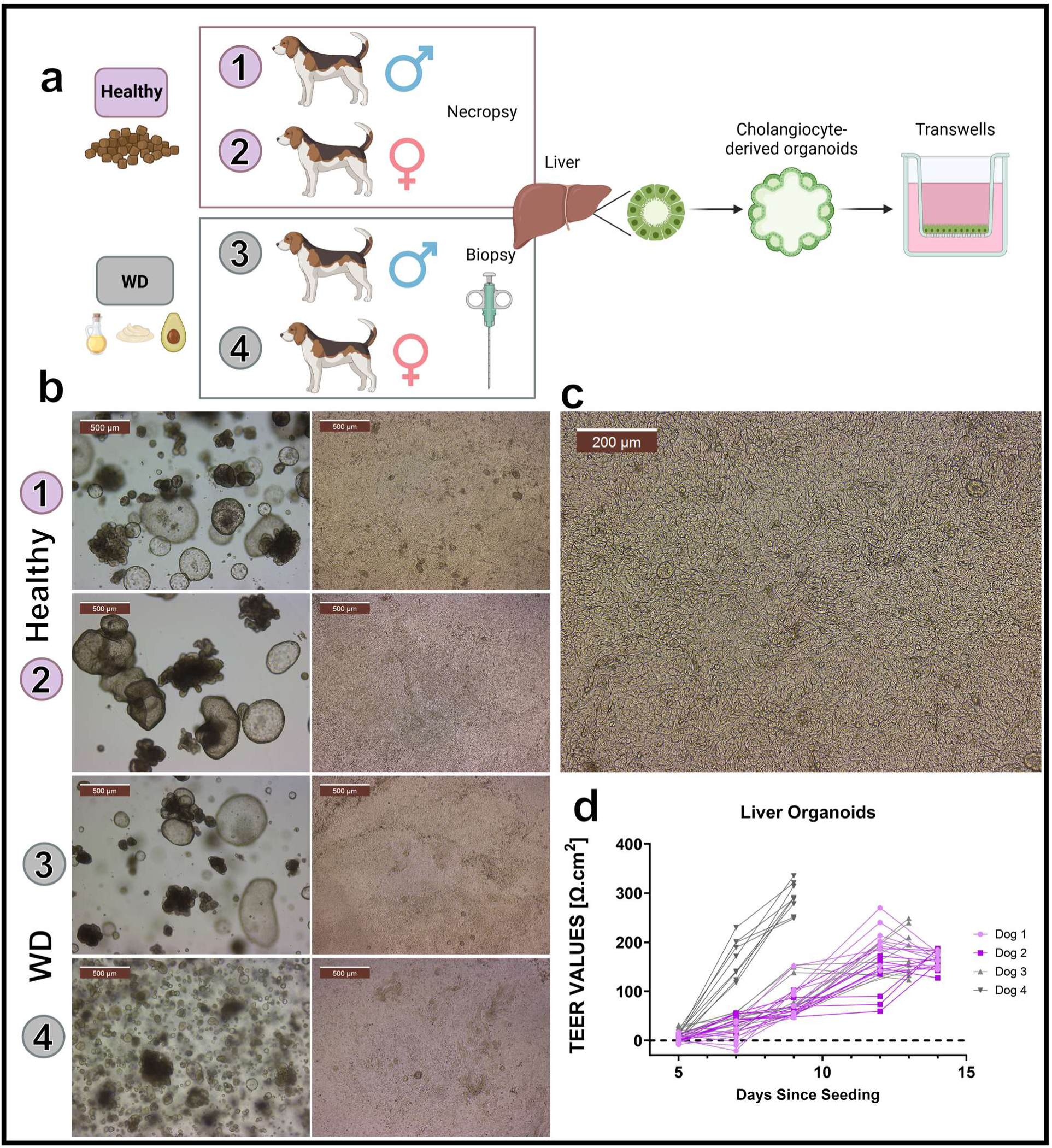
Establishment of canine liver organoid-derived polarized monolayers. (**a**) Experimental workflow. Representative brightfield microscopy images of (**b**) three-dimensional canine liver organoids and organoid-derived Transwell monolayers established from each donor. Organoid images were acquired on the day of dissociation into single cells, whereas monolayer images show untreated control wells on the final day of the experiment. (**c**) Higher-magnification image of a confluent canine organoid-derived cholangiocyte monolayer. Scale bars are indicated in µm. (**d**) Transepithelial electrical resistance (TEER) measurements for all samples over the full culture period. Dapagliflozin and bexagliflozin were added 24 h before the final TEER measurement.

### Histological and microscopic appearance

Using transmission electron microscopy, we observed clear apical-basal polarity in the monolayers and identified apical microvilli (**Figure 2a**). Ultrastructural hallmarks of epithelial differentiation, including tight junctions, were also evident across samples. H&E staining showed no qualitative differences in monolayer morphology between donors. Across donors, monolayers consisted predominantly of low columnar to cuboidal-flattened cells (**Figure 2b**), morphologically consistent with a cholangiocyte-enriched epithelium. PAS staining revealed qualitatively higher cytoplasmic glycogen in WD-derived monolayers than in healthy controls, and diastase digestion confirmed the staining was indeed glycogen-specific (**Figure 2c-d**).

**Figure 2:**
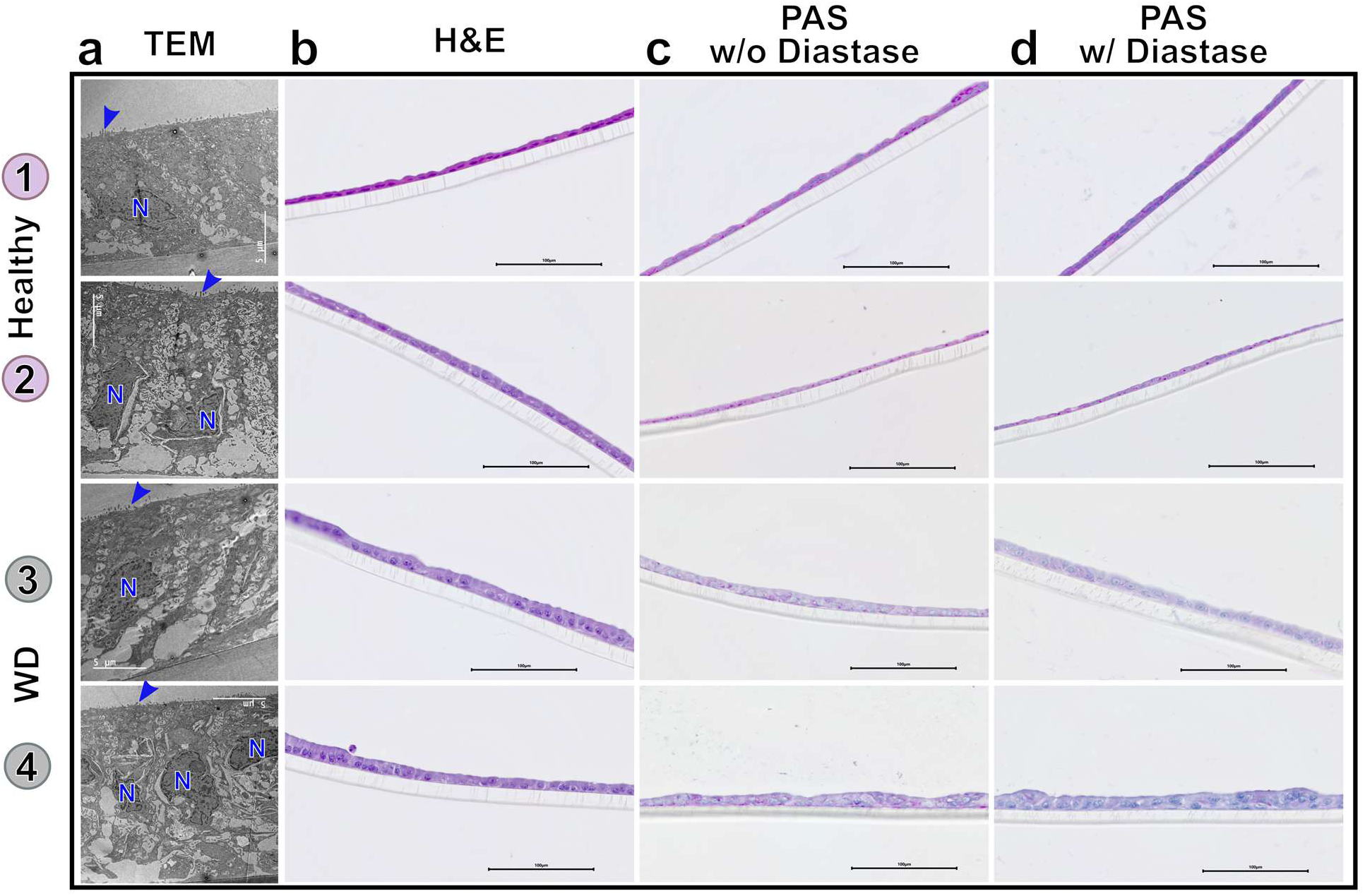
Histological characterisation of canine liver organoid–derived monolayers. (**a**) Transmission electron microscopy (TEM) of canine liver organoid–derived monolayers. Microvilli are shown with blue arrows while the nucleus is denoted with an “N”. (**b**) Haematoxylin and eosin staining of vehicle-control monolayers from healthy and WD donors, showing a uniform low columnar to cuboidal-flattened epithelium across donors with no overt treatment-associated differences. (**c**) Periodic acid-Schiff (PAS) staining showing qualitatively greater glycogen accumulation in WD-derived monolayers. (**d**) Diastase digestion (right column) diminishes the magenta PAS signal, confirming glycogen specificity. Scale bars, 100 µm.

### Transcriptomic expression

Bulk RNA-sequencing showed that organoid-derived Transwell monolayers exhibited a mature cholangiocyte phenotype, characterized by expression of *KRT18*, *PIGR*, *ANXA4*, and *SPP1*, while lacking canonical hepatocyte markers (**Figure 3e**) (Andrews et al., 2022). Additionally, there was low to absent expression of both *SLC5A1* and *SLC5A2* in the organoid models.

**Figure 3:**
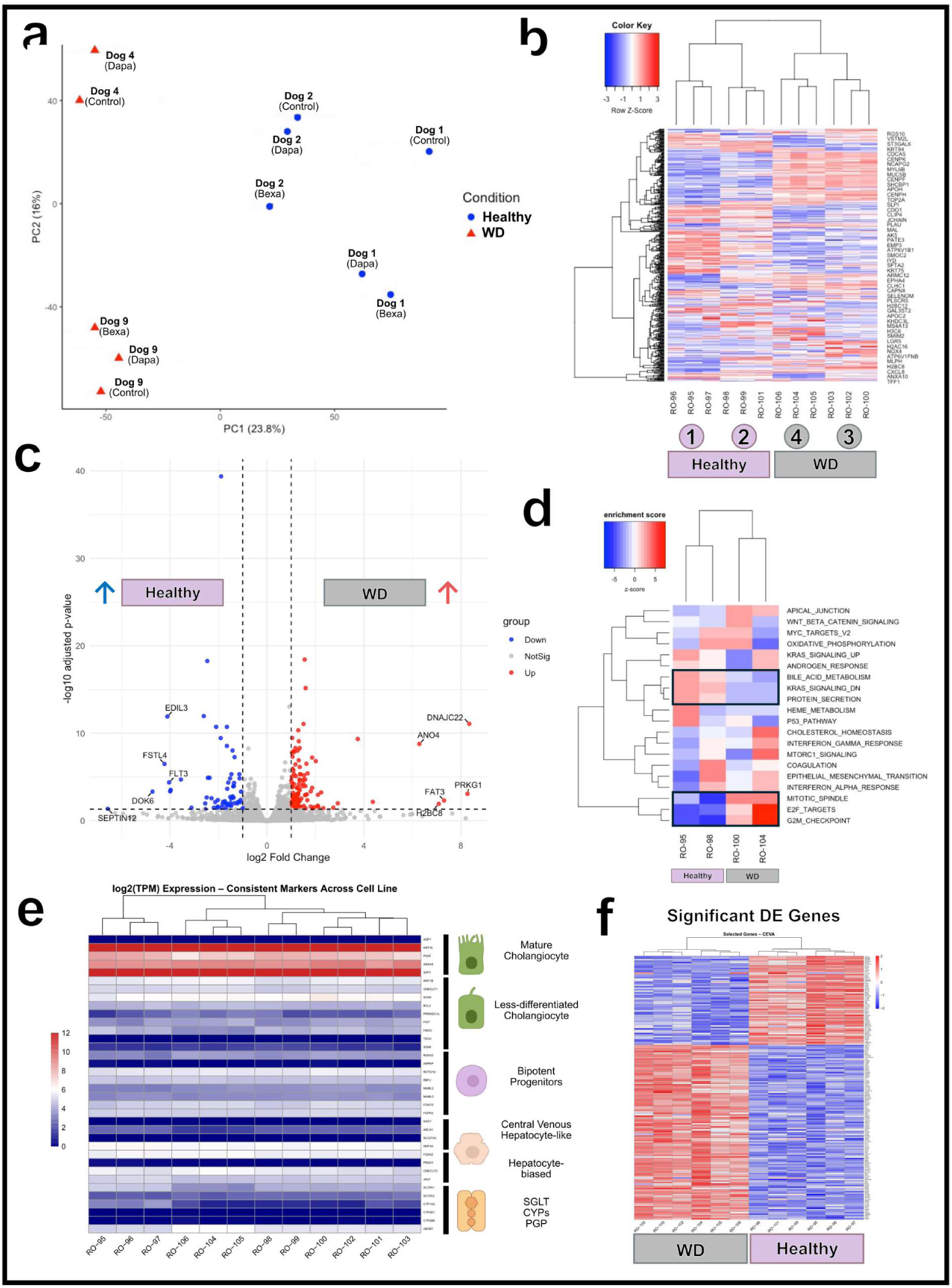
Transcriptomic profiling of canine liver organoids by health status. (**a**) PCA scores plot showing separation of healthy and WD-derived organoid samples on PC1. (**b**) Heatmap of the top 500 differentially expressed genes between healthy and WD organoids; rows clustered by Euclidean distance, columns by donor. (**c**) Volcano plot of differential expression (log_2_ fold change vs. −log_10_ adjusted P), highlighting transcripts upregulated in healthy (left) and WD (right). (**d**) Top pathway enrichments for the WD versus healthy contrast. (**e**) Cell type markers spanning cholangiocytes, progenitors, and hepatocytes, with relative expression shown as transcripts per million (TPM) (Andrews et al., 2022). (**f**) Differentially expressed genes identified by diet comparison, including all significantly upregulated and downregulated genes.

Transcriptomic differences were observed between monolayers derived from healthy and WD liver organoids (**Figure 3a-c**). Genes upregulated in WD-derived monolayers were markers associated with proliferation (*TOP2A, MKI67*) and fibrosis/extracellular matrix reprogramming (*COL4A1, COL4A2*) (**Figure 3f**). Consistent with these findings, pathway analysis identified significant upregulation of cell cycle-related programs, including *E2F* targets (**Figure 3d**). By contrast, significant transcriptomic differences were not detected between vehicle, dapagliflozin, and bexagliflozin within any donor or across donors.

### General metabolomics

The aligned general metabolomics peak table included 756 variables, and 47 observations after controls were dropped. One apical sample was removed from the dataset due to a failed GC injection. These samples were projected into PCA space (**Figure 4a**), where an obvious separation along principal component (PC) 2 between apical and basal supernatant was observed. Because the dominant axis of variation separated the apical from the basal compartment rather than the biological factors of interest (drug treatment and donor diet), apical and basal data were analyzed independently.

**Figure 4:**
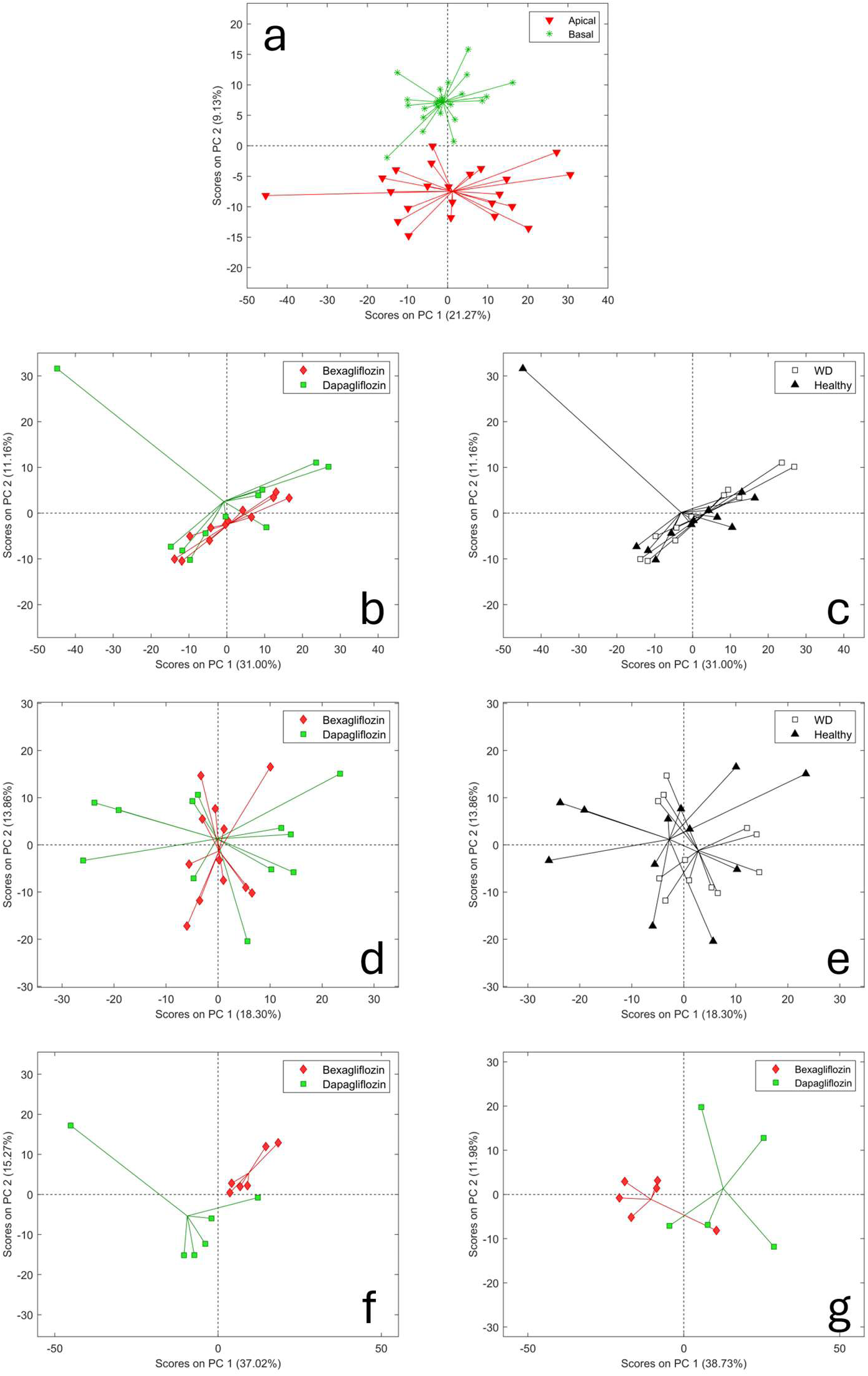
**(a)** PCA scores plot of all general metabolomics samples before feature selection, based on 756 detected metabolites and excluding controls. Samples are labeled according to compartment identity (apical or basal), revealing clear separation along PC2. **(b)** PCA scores plot of apical samples from healthy and Western diet (WD)-derived monolayers treated with either bexagliflozin or dapagliflozin. **(c)** PCA scores plot of apical samples treated with bexagliflozin or dapagliflozin, grouped by donor diet status (healthy or WD). **(d)** PCA scores plot of basal samples from healthy and WD-derived monolayers treated with either bexagliflozin or dapagliflozin. **(e)** PCA scores plot of basal samples treated with bexagliflozin or dapagliflozin, grouped by donor diet status (healthy or WD). **(f)** PCA scores plot of apical healthy state samples treated with either bexagliflozin or dapagliflozin. **(g)** PCA scores plots of apical WD samples treated with either bexagliflozin or dapagliflozin.

When apical samples were tested with health status and SGLT2 inhibitor as combined factors in a single-factor PERMANOVA model considering all 756 metabolites, neither term reached statistical significance, and PCA showed only limited separation in the corresponding score space (**Supporting Table 1**). The same was true of basal samples. Disaggregating the analysis by health state then revealed the apical-compartment drug separations described below; we caution that this stratification increases the number of pairwise tests and that any nominal significance should be interpreted in that light.

Two health-state-stratified apical comparisons reached nominal PERMANOVA significance prior to feature selection: the healthy-apical drug comparison (*P* = 0.0087, pseudo-F = 2.52, R^2^ = 0.202; **Figure 4f**) and the WD-apical drug comparison (*P* = 0.0149, pseudo-F = 2.73, R^2^ = 0.233; **Figure 4g**). For both comparisons, PERMDISP was not significant (**Supporting Table 3**), suggesting that these results were not attributable to differences within group dispersion (Anderson, 2006). FS-CR was then applied to identify the metabolites driving this separation. For healthy apical samples, 19 metabolites were identified as significant contributors to group separation, with 6 positively correlated with dapagliflozin and 13 with bexagliflozin, or which six were putatively identified (**Table 2**). For WD apical samples, 12 metabolites were identified as significant, all of which were associated with dapagliflozin treatment, with only four metabolites putatively identified: 2,3-dihydroxypyridine, 2,3-dihydroxybutanoic acid, 1-monostearin, and 1-methylinosine. Notably, there was no overlap in significant metabolites between healthy and WD apical sample comparisons.

**Table 2:** Features retained by FS-CR as contributing to the apical-compartment separation between dapagliflozin and bexagliflozin in healthy-donor monolayers (general metabolomics platform) with an MSI Level 2 putative identification. Italicised entries are positively correlated with dapagliflozin; non-italicised entries are positively correlated with bexagliflozin.

| **Metabolite** |
|---|
| *N-acetylmannosamine* |
| succinic acid |
| serine |
| homoserine |
| 2-picolinic acid |
| 2-deoxyerythritol |

We initially considered whether the apparent apical-compartment separations might reflect differential drug biotransformation rather than divergent mechanisms of action. To investigate this hypothesis, custom scripts were used to screen the raw data for chlorinated metabolites, which would indicate the presence of bexagliflozin or dapagliflozin metabolites not captured in standard mass spectral libraries. However, no chlorinated metabolites were putatively identified corresponding to potential or known metabolites of either drug. Metabolite 345401, overrepresented in dapagliflozin samples, was identified as containing one chlorine atom (**Supporting Figure 1**), however, it does not appear to be a metabolite of dapagliflozin. Because the metabolite sets differentiating dapagliflozin and bexagliflozin did not overlap between healthy and WD organoids, and no signals were conserved across omics platforms, these patterns likely reflect subtle, background-dependent metabolic variation rather than discrete drug mechanisms of action.

The following metabolites from four pathways, tricarboxylic acid (TCA) cycle, glycolysis, amino acid synthesis, and cholesterol were putatively identified; α-ketoglutarate, glucose, glucose-6-phosphate, alanine, leucine, isoleucine, valine, glycine, serine, cholesterol, and lithocholic acid (**Supporting Table 4**). No metabolites were statistically significant by *t*-test after FDR correction. Cholesterol’s Fisher ratio was notably large in apical healthy samples and apical WD samples when comparing SGLT2 inhibitors, with the abundance of cholesterol being higher in dapagliflozin cells in both cases (**Supporting Figures 2 & 3**). The inverse was true for lithocholic acid in apical healthy samples when comparing SGLT2 inhibitors (**Supporting Figure 4**).

### Lipidomics

The aligned lipidomics peak table included 3,069 variables, and 48 observations after removal of injection controls. Consistent with the general metabolomic findings, PCA revealed a dominant apical/basal separation, here along PC1, which accounted for 12.07 % of total variance (**Figure 5a**).

**Figure 5:**
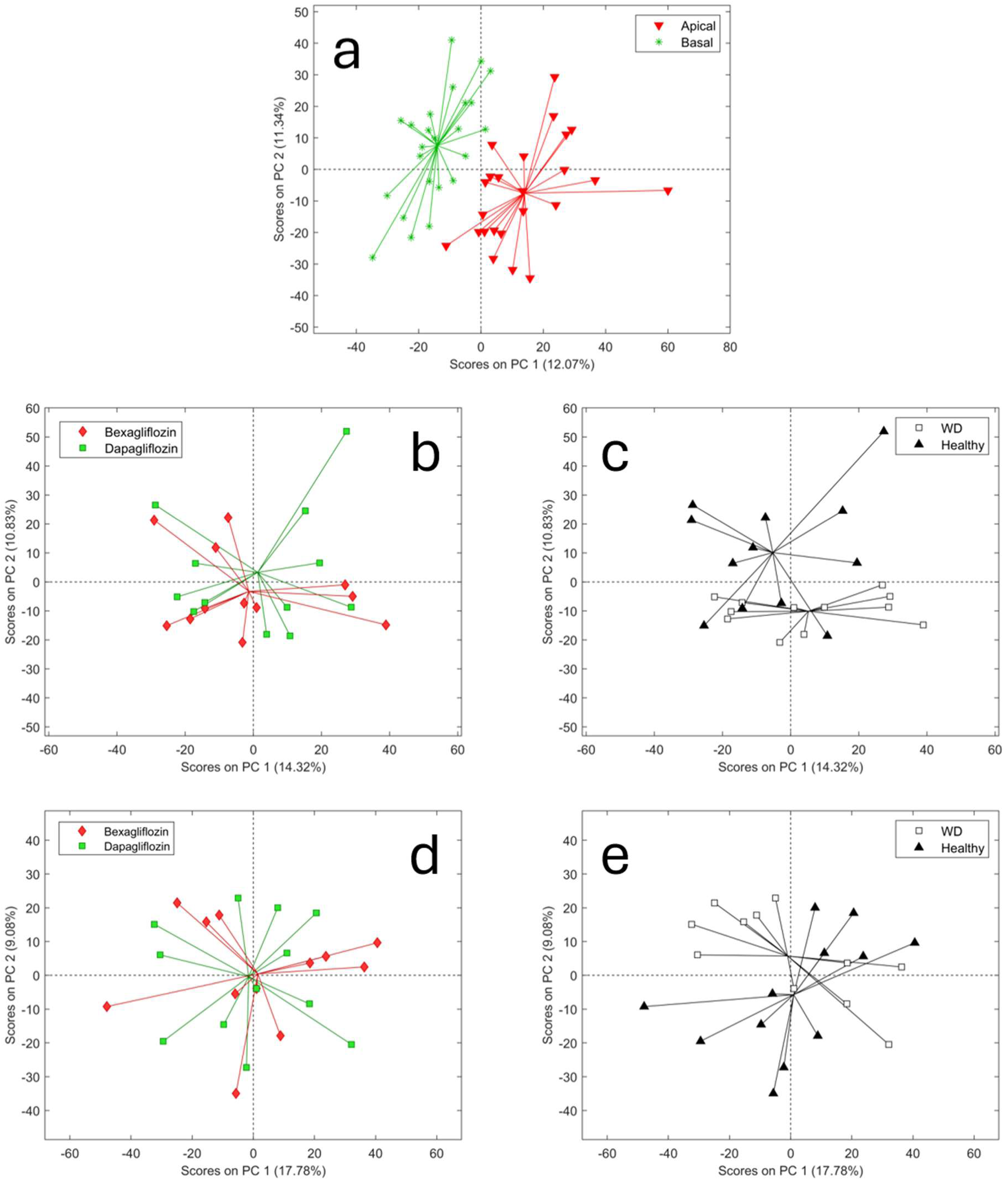
**(a)** PCA scores plot of all lipidomics samples before feature selection, based on 3,069 detected metabolites and excluding controls. Samples are labeled according to compartment identity (apical or basal), revealing clear separation along PC1. **(b)** PCA scores plot of apical samples from healthy and Western diet (WD)-derived monolayers treated with either bexagliflozin or dapagliflozin. **(c)** PCA scores plot of apical samples treated with bexagliflozin or dapagliflozin, grouped by donor diet status (healthy or WD). **(d)** PCA scores plot of basal samples from healthy and WD-derived monolayers treated with either bexagliflozin or dapagliflozin. **(e)** PCA scores plot of basal samples treated with bexagliflozin or dapagliflozin, grouped by donor diet status (healthy or WD).

Apical and basal lipidomic samples were analyzed independently. For apical samples, no statistically significant differences were observed between SGLT2 inhibitors by PERMANOVA when considering 3,069 variables before FS-CR (**Figure 5b**); however, a significant difference was found between healthy and WD samples (*P* = 0.0019, pseudo-F = 1.9051, R^2^ = 0.0797; **Figure 5c**). The PERMDISP was significant (**Supporting Table 3**) suggesting unequal within-group variability, suggesting the observed separation may reflect differences in centroids, dispersion, or both. Lipids positively associated with healthy samples in the apical compartment include the ceramides Cer 18:0;2O/24:1;O and Cer 18:1;2O/16:1, triglyceride TG 18:2_18:2_18:2;1O, and lysophosphatidylcholines LPC 20:2, LPC O-18:0, and LPC 18:0 (**Table 3**). The remaining detected lipids were not putatively identified to MSI Level 2.

**Table 3:**
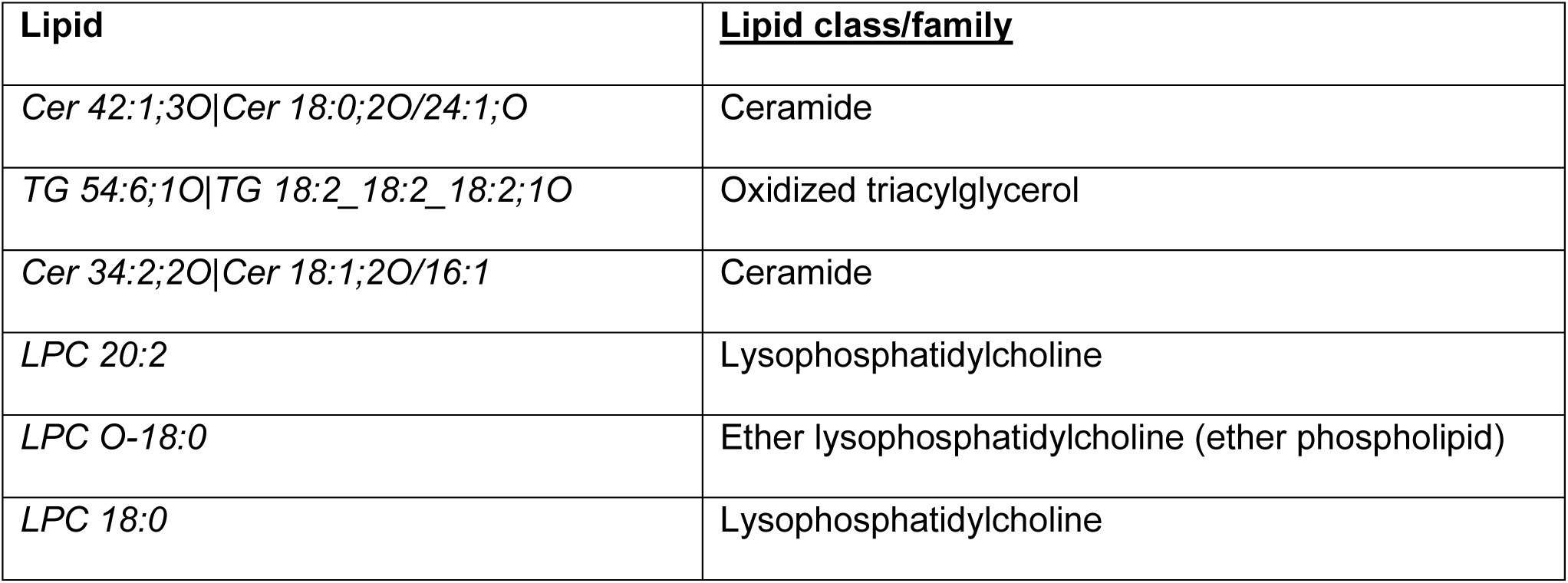
Features retained by FS-CR as contributing to the apical-compartment separation between healthy and WD samples exposed to either bexagliflozin or dapagliflozin (lipidomics platform) with MSI Level 2 putative identification. All lipids are positively correlated with healthy status. Lipid class/family are provided for each putatively identified lipid.

| **Lipid** | **Lipid class/family** |
|---|---|
| *Cer 42:1;3O\|Cer 18:0;2O/24:1;O* | Ceramide |
| *TG 54:6;1O\|TG 18:2_18:2_18:2;1O* | Oxidized triacylglycerol |
| *Cer 34:2;2O\|Cer 18:1;2O/16:1* | Ceramide |
| *LPC 20:2* | Lysophosphatidylcholine |
| *LPC O-18:0* | Ether lysophosphatidylcholine (ether phospholipid) |
| *LPC 18:0* | Lysophosphatidylcholine |

The same pattern was observed in the basal compartment: no significant differences were detected between SGLT2 inhibitor treatments before FS-CR (**Figure 5d**), whereas healthy and WD-derived samples differed significantly (*P* = 0.0324, pseudo-F = 1.5345, R^2^ = 0.0652; **Figure 5e**). This comparison was not significant by PERMDISP (**Supporting Table 3**), suggesting this result was not attributable to differences within group dispersion. Lipids positively associated with healthy samples in the basal compartment include the fatty acids palmitoleic acid and physeteric acid, triglycerides TG 49:1, TG 15:0_16:0_16:0, and TG 60:3, hexosylceramide HexCer 18:1;2O/16:0;O, and lysophosphatidylcholine LPC O-18:0 (**Table 4**). The lysophosphatidylcholine LPC 20:4 and ceramide Cer 18:0;2O/17:0;O were associated with WD (**Table 4**).

**Table 4:** Features retained by FS-CR as contributing to the basal-compartment separation between healthy and WD samples exposed to either bexagliflozin or dapagliflozin (lipidomics platform) with MSI Level 2 putative identification. Italicised entries are positively correlated with healthy status; non-italicised entries are positively correlated with WD. Lipid class/family are provided for each putatively identified lipid.

| **Lipid** | **Lipid class/family** |
|---|---|
| *CAR 14:2* | Acylcarnitine |
| *FA 16:1 (palmitoleic acid)* | Free fatty acid |
| *HexCer 34:1;3O\|HexCer 18:1;2O/16:0;O* | Hexosylceramide |
| *TG 49:1* | Triacylglycerol |
| *FA 14:1 (physeteric acid)* | Free fatty acid |
| *TG 47:0\|TG 15:0_16:0_16:0* | Triacylglycerol |
| *TG 60:3* | Triacylglycerol |
| *Cer d36:1* | Ceramide |
| *LPC O-18:0* | Ether lysophosphatidylcholine |
| LPC 20:4 | Lysophosphatidylcholine |
| Cer 35:0;3O\|Cer 18:0;2O/17:0;O | Ceramide |

Effect sizes are modest in absolute terms but consistent across compartments, reinforcing the diet-driven nature of the lipidomic signal.

### Biogenic amines

The aligned biogenic amines peak table comprised 3,406 variables, and 48 observations after removal of injection controls. Consistent with the patterns observed in both the general metabolomics and lipidomics datasets, PCA showed a clear apical/basal separation, most prominent along PC2, most likely attributed to the different volumes between apical and basal chambers (**Figure 6a**).

**Figure 6:**
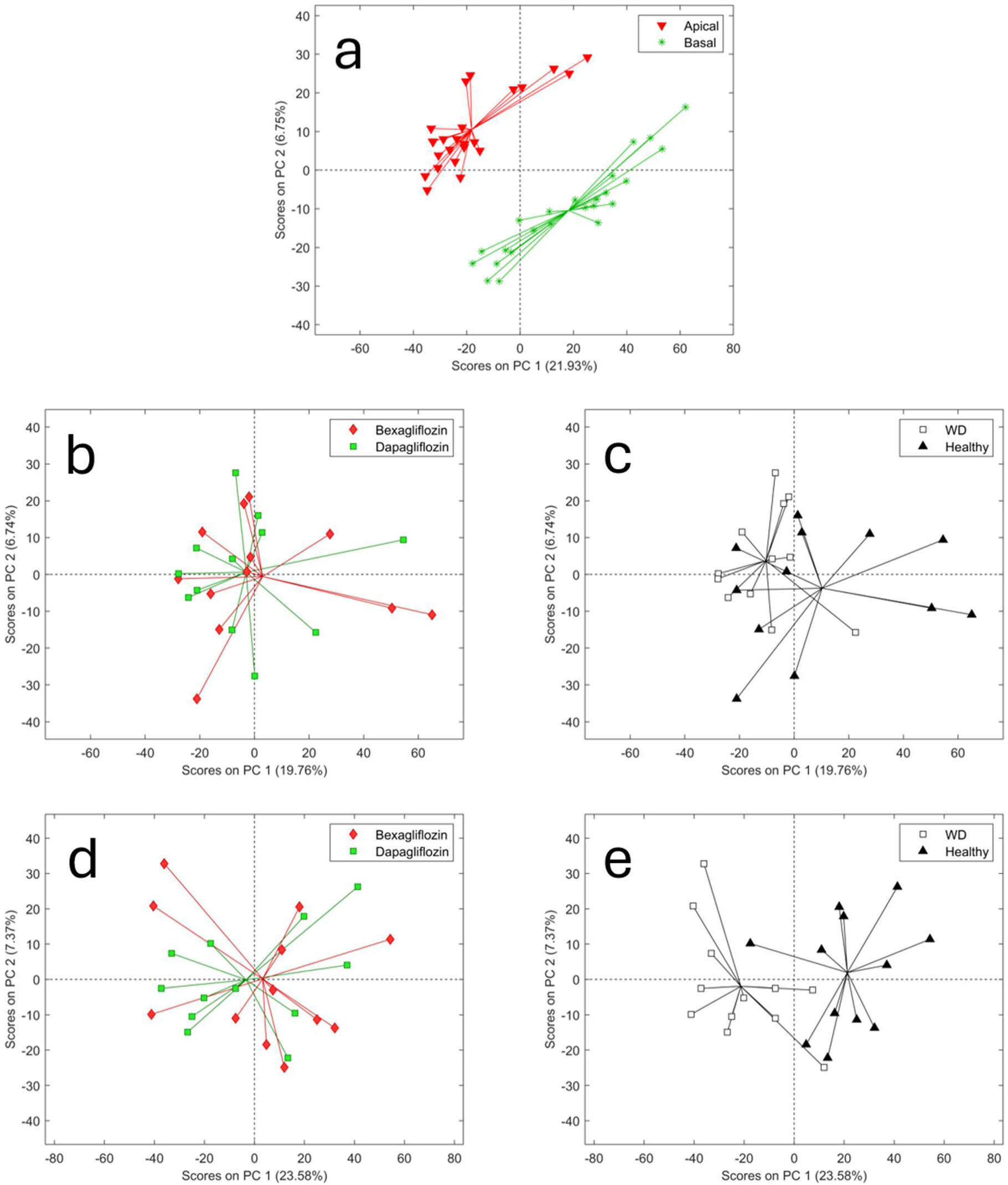
**(a)** PCA scores plot of all bioamine samples before feature selection, based on 3,406 detected metabolites and excluding controls. Samples are labeled according to compartment identity, apical or basal, revealing clear separation along PC2. **(b)** PCA scores plot of apical samples from healthy and Western diet (WD)-derived monolayers treated with either bexagliflozin or dapagliflozin. **(c)** PCA scores plot of apical samples treated with bexagliflozin or dapagliflozin, grouped by donor diet status, healthy or WD. **(d)** PCA scores plot of basal samples from healthy and WD-derived monolayers treated with either bexagliflozin or dapagliflozin. **(e)** PCA scores plot of basal samples treated with bexagliflozin or dapagliflozin, grouped by donor diet status, healthy or WD.

Apical and basal biogenic-amine samples were analyzed independently. In the apical compartment, no significant drug effect was detected before FS-CR (**Figure 6b**), but the healthy versus WD contrast reached statistical significance when considering all 3,406 variables (*P* = 0.0265, pseudo-F = 1.644, R^2^ = 0.070; **Figure 6c**). The PERMDISP was significant (**Supporting Table 3**) suggesting unequal within-group variability, suggesting the observed separation may reflect differences in centroids, dispersion, or both. Biogenic-amines positively associated with healthy samples include acetylspermine, pipecolic acid, 1-methylnicotinamide, and kynurenic acid (**Table 5**) while spermidine and 2-[(4-Aminobenzoyl)amino]acetic acid were positively associated with WD. Non-nitrogen containing, small polar metabolites such as citric acid and glycine were putatively identified (MSI Level 2).

**Table 5:**
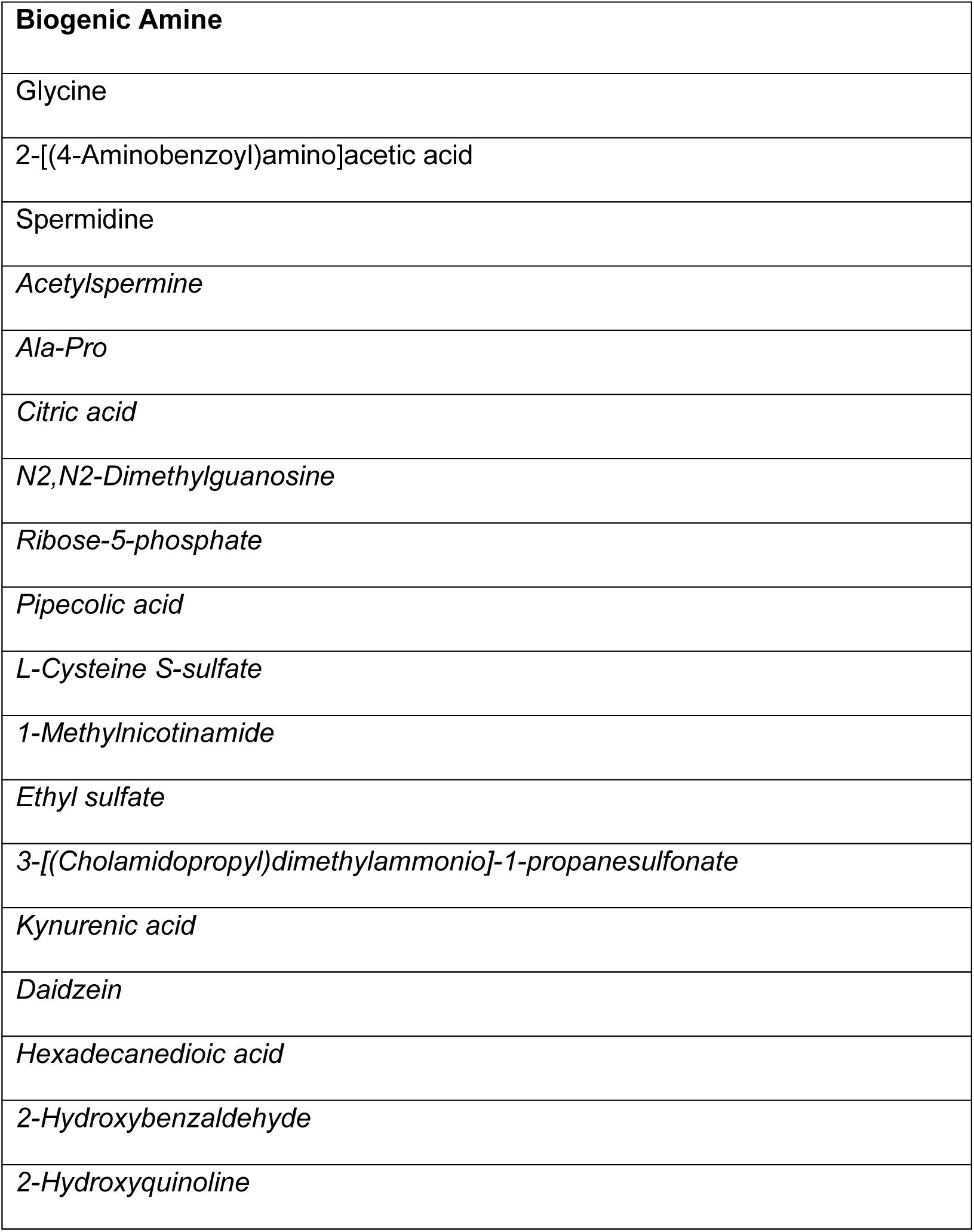
Features retained by FS-CR as contributing to the apical-compartment separation between healthy and WD samples exposed to either bexagliflozin or dapagliflozin (biogenic-amines platform) with MSI Level 2 putative identification. Italicised entries are positively correlated with healthy status; non-italicised entries are positively correlated with WD.

| **Biogenic Amine** |
|---|
| Glycine |
| 2-[(4-Aminobenzoyl)amino]acetic acid |
| Spermidine |
| *Acetylspermine* |
| *Ala-Pro* |
| *Citric acid* |
| *N2,N2-Dimethylguanosine* |
| *Ribose-5-phosphate* |
| *Pipecolic acid* |
| *L-Cysteine S-sulfate* |
| *1-Methylnicotinamide* |
| *Ethyl sulfate* |
| *3-[(Cholamidopropyl)dimethylammonio]-1-propanesulfonate* |
| *Kynurenic acid* |
| *Daidzein* |
| *Hexadecanedioic acid* |
| *2-Hydroxybenzaldehyde* |
| *2-Hydroxyquinoline* |

In the basal compartment, the drug effect was again non-significant (**Figure 6d**), while the health-status effect was the strongest observed in the entire study (P = 3 × 10^-5^, pseudo-F = 4.06, R^2^ = 0.156; **Figure 6e**). This comparison was not significant by PERMDISP (**Supporting Table 3**), suggesting this result was not attributable to differences within group dispersion. Biogenic-amines positively associated with healthy samples include uridine, N-Acetyl-D-galactosamine 4-sulfate, taurocholic acid, and tyrosine (**Table 6**), while only one unannotated metabolite was associated with WD. Similar to the annotations in **Table 5**, many non-amine metabolites were annotated in this assay; however, this is unsurprising as this HILIC-MS method cannot distinguish amine containing species from other small, polar, non-amine containing analytes. Given this, species such as 2-(N-Morpholino)ethanesulfonic acid (better known as MES buffer) and mannitol were detected and annotated.

**Table 6:** Features retained by FS-CR as contributing to the basal-compartment separation between healthy and WD samples exposed to either bexagliflozin or dapagliflozin (biogenic-amines platform) with MSI Level 2 putative identification. All features are positively correlated with WD.

| **Biogenic Amine** |
|---|
| 2-(N-Morpholino)ethanesulfonic acid |
| Gln-Phe |
| Uridine |
| 2,8-Quinolinediol |
| N-Acetyl-D-galactosamine 4-sulfate |
| allo-Inositol |
| Taurocholic acid |
| Sorbose |
| Acetylenedicarboxylic acid |
| Mannitol |
| Mannose |
| Tyrosine |

## DISCUSSION

The principal contribution of this study is the demonstration that polarized canine liver organoid-derived monolayers retain a measurable subset of the *in vivo* WD-associated phenotype after expansion, cryopreservation, dissociation, and growth on Transwell membranes. Several independent findings from the present study suggest that canine liver organoid-derived polarized monolayers retain selected features of *in vivo* WD-associated hepatic reprogramming. Donor-of-origin health status is at least partially retained after expansion, as evidenced by concordant healthy-versus-WD separation across transcriptomics, lipidomics, biogenic amines, and PAS histology. The system preserves apical-basal polarity, enabling compartment-resolved sampling that approximates the canalicular-sinusoidal organization of the *in vivo* hepatobiliary unit. The transcriptomic WD signature shows partial qualitative concordance with the *in vivo* phenotype reported in the parent cohort of WD-fed dogs (Mason et al., 2025), supporting back-translation from the *in vitro* platform to the *in vivo* donor.

Diet-associated differences were detected in lipidomic and biogenic amine profiles, in bulk transcriptomic separation, and qualitatively in PAS-positive glycogen accumulation in WD-derived monolayers. However, the general polar metabolome, the platform most directly reflecting glycolytic, tricarboxylic acid (TCA) cycle, and free amino acid pools, did not show a statistically significant diet effect in either compartment. PERMANOVA R^2^ values for the diet contrast were 0.080 in apical lipidomics, 0.065 in basal lipidomics, 0.070 in apical biogenic amines, and 0.156 in basal biogenic amines, with the basal biogenic amine compartment yielding the smallest P value across the dataset. These effect sizes were modest in absolute terms, indicating that diet explained only a minority of the variance in each metabolomic class. Nevertheless, the detection of diet-associated differences across lipidomic, biogenic amine, transcriptomic, and histologic readouts indicates that the model preserves aspects of the WD-associated epithelial phenotype. In particular, the WD-associated transcriptomic signature, comprising upregulation of proliferation markers (*TOP2A*, *MKI67*), extracellular matrix-associated genes (*COL4A1*, *COL4A2*), and enrichment of E2F target genes, resembles transcriptional features reported during cholangiocyte activation and ductular reaction in steatohepatitis, which correlate with fibrosis severity in human metabolic dysfunction-associated steatotic liver disease (Gadd et al., 2014; Sato et al., 2019), and are induced by WD feeding in rodent models of steatohepatitis (Yang et al., 2020). Given the exploratory design and the limited biological replication (two donors per group), these observations should be interpreted as hypothesis-generating and will require confirmation in a larger cohort. The absence of a comparable diet effect in the general polar metabolome may reflect several non-mutually exclusive mechanisms discussed below.

Two features of the general-metabolomics analysis deserve specific comment. First, the unsupervised variance structure was dominated by an apical-basal compartment separation, driven by differences in volumes between compartments, compartment-specific secretion, and uptake across a polarized epithelium. Second, and more importantly, neither drug treatment nor health status reached statistical significance when both factors were tested in the same PERMANOVA model in either compartment. We considered whether differential drug biotransformation could explain the apical-compartment separations. Both dapagliflozin and bexagliflozin are cleared predominantly by hepatic oxidation and glucuronidation, and the chlorine moiety present in each parent compound is retained in known phase I and phase II metabolites (Gumieniczek & Berecka-Rycerz, 2023). We therefore screened the GC-MS dataset for chlorinated features (Nam et al., 2021) but did not identify features consistent with reported metabolites of either compound. The only chlorine-containing feature retained by FS-CR, metabolite 345401, which was overrepresented in dapagliflozin-treated apical samples, had a mass spectrum inconsistent with documented dapagliflozin metabolites. This analysis argues against detectable phase I biotransformation products as the primary driver of the apical separation between SGLT2 inhibitors, but it does not exclude drug metabolism more broadly. GC-MS has limited sensitivity for glucuronide conjugates, which represent a major metabolic pathway for both compounds; targeted LC-MS analysis of expected phase II metabolites would therefore be required to formally exclude biotransformation as a contributor. Accordingly, we conclude only that the available GC-MS data do not support biotransformation as the main explanation for the apical signal. The absence of drug-associated separation in lipidomics, biogenic amines, and transcriptomics, together with the non-overlap between the healthy-apical and WD-apical FS-CR feature lists and the limited biological replication of the study design, precludes a mechanistic interpretation of the drug-associated apical separations in this dataset. We therefore interpret these apical drug-associated separations as exploratory and more likely attributable to donor-level variability than to a reproducible drug-specific metabolic response.

The absence of a diet effect on this platform is itself informative (**Supporting Table 1**). The polar metabolome captured by GC-TOF is weighted toward glycolytic intermediates, TCA-cycle metabolites, free amino acids, and small organic acids; exactly the pools we would expect to be remodeled by chronic WD feeding *in vivo*, and which the transcriptomic and PAS-histological data here suggest are at least partially remodeled in the WD donors. The fact that this signal does not persist in the *in vitro* Transwell platform may reflect the cholangiocyte-enriched nature of the monolayers, which may not capture hepatocyte-dependent components of the WD-associated metabolic phenotype. In addition, analysis of the culture supernatant may be insufficient to resolve diet-driven changes in intracellular central carbon metabolism, particularly if these changes are more readily reflected in lipidomic or biogenic amine outputs. PAS-histologic data also represents a single snapshot in time of a limited subset of the cells on the Transwells and may not be representative of the cellular metabolic state. Finally, a smaller diet-associated effect in the polar metabolome may remain below the detection threshold of the present experimental design.

A targeted pathway-level analysis identified one consistent qualitative pattern: cholesterol abundance was higher in dapagliflozin-treated apical samples in both healthy and WD donors, with Fisher ratios of 4.18 and 10.69, respectively, whereas lithocholic acid showed the opposite pattern in healthy apical samples. Neither contrast survived FDR correction across the full untargeted matrix, and both signals were restricted to the apical compartment, suggesting that they may reflect compartment-specific secretion rather than intracellular metabolic reprogramming. We therefore interpret these findings as exploratory/hypothesis-generating only. The parallel to the reported clinical effect of SGLT2 inhibitors, a modest rise in circulating LDL and total cholesterol, is directional only. Basu et al. linked the increase in LDL in a diabetic mouse model to reduced LDL clearance and enhanced lipolysis of triglyceride-rich lipoproteins, mechanisms that require an intact circulating lipoprotein system and cannot be reproduced in an isolated, cholangiocyte-enriched Transwell monolayer without functional hepatocyte and vascular compartments (Basu et al., 2018). The cholesterol signal observed here must therefore arise from processes intrinsic to the culture system, including altered uptake, intracellular handling, membrane turnover, or compartmental transport, rather than from systemic LDL metabolism. Moreover, the dominant hepatic pathways governing circulating lipoprotein metabolism and classical CYP7A1-mediated bile-acid synthesis are hepatocyte-centered and underrepresented in this model. A transport-level explanation is therefore more compatible with the platform. Cholangiocytes actively modify hepatocyte-derived bile through both absorptive and secretory processes (Tabibian et al., 2013) and express the apical bile-acid transporter ASBT, which mediates bile-acid uptake (Alpini et al., 2005). Thus, the apical increase in cholesterol accompanied by a decrease in lithocholic acid may reflect altered polarized transport or transcellular handling rather than increased net synthesis. This interpretation remains speculative because lithocholic acid is generated primarily through microbial conversion of primary bile acids, making its putative identification in a microbiome-free monolayer uncertain (Ridlon et al., 2006). Mechanistic interpretation would require absolute quantification using authentic standards, isotope-labeled flux tracing, evaluation across a clinically relevant concentration range, and biological replication beyond the two donors per group included in this study.

Lipidomic profiles recapitulated the apical and basal findings observed in the general metabolomics dataset and showed no drug-associated separation in either compartment. In contrast, a statistically significant healthy-versus-WD difference was detected in both apical and basal samples. The corresponding effect sizes were modest in absolute terms, with diet accounting for approximately 7% of total variance in the lipidomic feature space. Nevertheless, the direction of the diet-associated separation was consistent across compartments and concordant with the transcriptomic separation between healthy and WD-derived monolayers, as well as with the increased PAS-positive glycogen accumulation observed in WD-derived monolayers (**Table 3**; **Table 4**). LPC O-18:0, an ether-linked phospholipid, was associated with healthy cells in both apical and basal compartments. Ether-linked phospholipids have been associated with protecting against oxidative stress through their antioxidant properties (Dean & Lodhi, 2018). This is potentially indicative of reduced oxidative stress relative to WD cells. However, the limited annotation of the lipidome, particularly discriminating lipid features, presents a significant bottleneck in interpreting the observed lipidome-wide reprogramming (Sumner et al., 2007). The untargeted lipidomics results here largely suggest that triacylglycerol, ceramides, free fatty acids, and lysophospholipids are driving the separation in health-status, therefore future work should use targeted lipidomics to identify and quantify lipids in these lipid families in a larger replication study.

Biogenic amine profiles followed the platform-wide apical-basal separation pattern, with no detectable drug-associated effect in either compartment and a statistically significant healthy-versus-WD contrast in both. The apical contrast was modest, whereas the basal contrast represented the largest health-status effect observed across the dataset. The basal biogenic amine R^2^ value of 0.156 was notably higher than the 0.065-0.080 range observed for lipidomics and apical biogenic amines, and this contrast is therefore the most likely to remain significant under a donor-stratified permutation re-analysis. As many metabolites detected in the biogenic amine assay were not putatively identified to MSI level 2, the most direct next experiment will be to identify the specific features driving this separation and to perform absolute quantification of canonical biogenic amine pathway metabolites, including urea-cycle intermediates, polyamines, aromatic amino acids and their derivatives, and methylated amines, in a larger replication cohort.

Several limitations warrant explicit acknowledgement. First, biological replication is limited to two donors per health state and per drug arm. Second, the monolayers are cholangiocyte-enriched rather than hepatocyte-mature, which materially restricts the hepatic processes that can be interrogated with this model and likely contributes to the absence of a diet-associated effect in the general metabolome. In particular, cholangiocytes are not the principal cellular compartment for glycolytic, TCA-cycle, and broader hepatocellular metabolic reprogramming. Future work should therefore prioritize increased biological replication across additional donors; donor-stratified re-analysis of the present dataset; targeted, absolutely quantified metabolomics of bile acid, cholesterol, urea-cycle, polyamine, and aromatic amino acid pathways; and isotope-labeled glucose, palmitate, and glutamine flux analyses to test the candidate cholesterol and bile acid signals. Future efforts to differentiate, or co-culture the cholangiocyte organoids with hepatocytes would greatly extend the usefulness of the model. Third, the healthy and Western-diet lines differed in provenance, sampling modality, and husbandry as well as in dietary exposure. The two healthy donors were commercial-vendor dogs (Marshall BioResources) sampled post-mortem at necropsy and maintained on a standard grain-based maintenance kibble, whereas the two Western-diet donors were research-colony beagles sampled by laparoscopic biopsy after eight weeks on a WD described previously by our consortium (Mason et al., 2025). Gonadal status was moreover confounded with diet group: both Western-diet donors were surgically sterilized (one neutered male, one spayed female), whereas both healthy donors were sexually intact, so any sex-hormone-dependent contribution to the observed differences cannot be separated from the dietary effect. The healthy-versus-WD contrast therefore conflates the dietary intervention with differences in donor source, husbandry, reproductive status, and tissue-collection method, and in practical terms compares a standard maintenance kibble against a formulated WD. Diet-attributable interpretations of the observed signatures should accordingly be read with this confound in mind, and the earlier back-translation and cross-platform concordance statements should be interpreted in the same light. Overall, this study design was effective for hypothesis generation, allowing us to examine the effects of WD exposure and drug treatment across three complementary analytical platforms. While broad metabolite coverage was achieved, the low rate of putative metabolite identification hinders biological interpretation. Future studies could improve metabolite annotation through isotope labeling and flux analysis, while targeted metabolomics would enable deeper interrogation of specific pathways of interest. This is particularly relevant for the pathways examined here through manual extraction of putatively identified metabolites. Several pathway-associated metabolites showed nominal statistical differences; however, these associations did not remain significant after FDR correction across the full metabolite matrix. Targeted, quantitative metabolomic analyses will therefore be needed to evaluate these candidate pathways more rigorously and to test the hypotheses generated from this exploratory dataset. It is also important to note that the Transwell system contained different volumes in the apical (200 μL) and basal (700 μL) chambers, and this volume difference alone may be the main cause for separation of apical and basal samples. Given this, most analysis focused on inter-apical and inter-basal comparisons.

## CONCLUSION

This study demonstrates that canine liver organoid-derived monolayers may recapitulate WD-induced hepatic reprogramming at the transcriptomic, lipidomic, and biogenic-amine levels, with corroborating qualitative histological evidence of increased PAS-positive, diastase-sensitive material consistent with glycogen accumulation in WD donors. We found dapagliflozin and bexagliflozin produced only subtle, context-dependent metabolic perturbations.

## Supporting information

Full Summary

## Availability of data and material

The stranded mRNA raw RNA-seq reads generated and analyzed in this study are available in the NCBI GEO. RNA-seq scripts are available on GitHub (https://github.com/chris-zdyrski/Liver_Organoids). Raw metabolomics data (general metabolomics, biogenic amines, and lipidomics) are available on Borealis (https://doi.org/10.5683/SP4/KLXAXZ).

## Author Contributions

**R.T.G.:** Methodology, resources, formal analysis, investigation, data curation, visualization, writing – original draft, writing – review & editing. **C.Z.:** Conceptualization, methodology, validation, formal analysis, investigation, resources, data curation, visualization, writing – original draft, writing – review & editing. **P.d.l.M:** Methodology, visualization, writing – review & editing. **S.L.N.:** Methodology, visualization, writing – review & editing. **A.B.M.:** Conceptualization, investigation, writing – review & editing. **M.P.C.:** Methodology, formal analysis, investigation, writing – review & editing. **H.C.:** Formal analysis, investigation, visualization, writing – review & editing. **J.L.W.:** Investigation, writing – review & editing. **E.G.:** Resources, writing – review & editing. **E.D.:** Formal analysis, investigation, writing - review & editing. **J.J.H.:** Resources, visualization, writing – review & editing. **J.P.M.:** Conceptualization, methodology, validation, formal analysis, investigation, resources, data curation, visualization, supervision, project administration, funding acquisition, writing – original draft, writing – review & editing.

## Acknowledgments

We appreciate the timely processing of samples at the University of Georgia Histology Laboratory. We thank Tracey Stewart, manager of the Light and Electron Microscopy Roy J. Carver High-Resolution Microscopy Facility at Iowa State University, for her help and support throughout this project. We thank Abigail Ralston for her assistance with the organoid cultures.

## Declaration of Assistive AI in Scientific Writing

ChatGPT-5.5 (OpenAI, 2026), Grok 4.2 Beta (xAI, 2026), and Claude Opus 4.6/5 (Anthropic, 2026) were used to assist with the drafting and editing of this manuscript. Their use was exclusively limited to improving grammar and enhancing readability, without generating new data or analyses. Use of AI assistance is disclosed in line with prevailing authorship and publication-ethics guidance from the *Committee on Publication Ethics* (COPE^1^) and the *International Committee of Medical Journal Editors* (ICMJE^2^). All AI-generated text was reviewed and edited thoroughly by the authors, who take full responsibility for the final content of the publication.

## Footnotes

1 Committee on Publication Ethics. *COPE Core Practices*. COPE; 2017. Updated regularly. Available from: https://publicationethics.org

2 International Committee of Medical Journal Editors. *Recommendations for the Conduct, Reporting, Editing, and Publication of Scholarly Work in Medical Journals*. ICMJE; updated 2024. Available from: https://www.icmje.org

