## Supplementary material for "Polarized Canine Liver Organoid Monolayers Capture Features of Western Diet-Induced Hepatic Metabolic Reprogramming Across Transcriptomic, Lipidomic, and Biogenic Amine Signatures": SI

### Supporting File

The first tab, 'summary_all...' consists of a description of all comparisons, with all the models, and if they are significant or not by PERMANOVA. The next 4 are a tabulation of metabolites identified as significant by FS-CR for all comparisons (i.e.; all metabolites associated with healthy samples across all analytical platforms and comparisons). The remaining tabs contain the selected significant features from each comparison.

### Supporting Tables

*Supporting Table 1: Summary statistics from permutation multivariate analysis of variance (PERMANOVA) before feature selection for all 24 models. BA = bioamines, GM = general metabolomics, LP = lipidomics. When comparison reads 'BA Apical Drug', this is bioamines, apical compartment, comparing drugs, while including healthy and WD cells. 'BA Apical Healthy Drug' is bioamines, apical compartment, comparing drugs, but only differences between healthy cells.*

| Comparison | Significant | P value | F | $R^2$ |
|---|---|---|---|---|
| BA Apical Drug | No | 0.6758 | 0.8697 | 0.038 |
| **BA Apical Health** | **Yes** | **0.0265** | **1.6443** | **0.0695** |
| BA Apical Healthy Drug | No | 0.3085 | 1.0418 | 0.0944 |
| BA Apical WD Drug | No | 0.7102 | 0.9034 | 0.0829 |
| BA Basal Drug | No | 0.804 | 0.7866 | 0.0345 |
| **BA Basal Health** | **Yes** | **0.00003** | **4.064** | **0.1559** |
| BA Basal Healthy Drug | No | 0.9218 | 0.7972 | 0.0738 |
| BA Basal WD Drug | No | 0.6699 | 0.8816 | 0.081 |
| GM Apical Drug | No | 0.3257 | 1.026 | 0.0466 |
| GM Apical Health | No | 0.4266 | 0.9395 | 0.0428 |
| **GM Apical Healthy Drug** | **Yes** | **0.0087** | **2.5249** | **0.2016** |
| **GM Apical WD Drug** | **Yes** | **0.0149** | **2.734** | **0.233** |
| GM Basal Drug | No | 0.4255 | 0.9946 | 0.0433 |
| GM Basal Health | No | 0.179 | 1.2183 | 0.0525 |
| GM Basal Healthy Drug | No | 0.3166 | 1.1032 | 0.0994 |
| GM Basal WD Drug | No | 0.3454 | 1.0481 | 0.0949 |
| LP Apical Drug | No | 0.2258 | 1.109 | 0.048 |
| **LP Apical Health** | **Yes** | **0.0019** | **1.9051** | **0.0797** |
| LP Apical Healthy Drug | No | 0.0951 | 1.2705 | 0.1127 |
| LP Apical WD Drug | No | 0.1818 | 1.1593 | 0.1039 |
| LP Basal Drug | No | 0.3589 | 1.0143 | 0.0441 |
| **LP Basal Health** | **Yes** | **0.0324** | **1.5345** | **0.0652** |
| LP Basal Healthy Drug | No | 0.5092 | 0.9281 | 0.0849 |
| LP Basal WD Drug | No | 0.3611 | 1.0072 | 0.0915 |

*Supporting Table 2: Summary statistics from permutation multivariate analysis of variance (PERMANOVA) after feature selection for 11 models. BA = bioamines, GM = general metabolomics, LP = lipidomics. When comparison reads 'BA' Apical health, this is bioamines, apical compartment, comparing health status, while including both drugs.*

| Comparison | Significant | P value | F | $R^2$ |
|---|---|---|---|---|
| BA Apical Health | Yes | <0.00001 | 11.6577 | 0.3464 |
| BA Basal Health | Yes | <0.00001 | 29.1599 | 0.57 |
| GM Apical Health | Yes | <0.00001 | 10.3751 | 0.3307 |
| GM Basal Health | Yes | <0.00001 | 6.0579 | 0.2159 |
| LP Apical Health | Yes | <0.00001 | 17.0814 | 0.4371 |
| LP Basal Health | Yes | <0.00001 | 10.6801 | 0.3268 |
| BA Basal Drug | Yes | <0.00001 | 9.4916 | 0.3014 |
| GM Apical Drug | Yes | <0.00001 | 5.6353 | 0.2116 |
| GM Basal Drug | Yes | <0.00001 | 6.9063 | 0.239 |
| LP Apical Drug | Yes | <0.00001 | 17.5645 | 0.4439 |
| LP Basal Drug | Yes | <0.00001 | 12.5064 | 0.3624 |

*Supporting Table 3: Summary statistics from Permutational Analysis of Multivariate Dispersions (PERMDISP) before feature selection for all models significant by PERMANOVA (Supporting Table 1). BA = bioamines, GM = general metabolomics, LP = lipidomics. When comparison reads 'BA Apical Drug', this is bioamines, apical compartment, comparing drugs, while including healthy and high fat diet cells. 'BA Apical Healthy Drug' is bioamines, apical compartment, comparing drugs, but only differences between healthy cells.*

| Comparison | Significant | P value | F |
|---|---|---|---|
| BA Apical Health | Yes | 0.00719 | 8.5682 |
| BA Basal Health | No | 0.0642 | 3.7806 |
| GM Apical Healthy Drug | No | 0.2275 | 1.6411 |
| GM Apical WD Drug | No | 0.1522 | 2.0955 |
| LP Apical Health | Yes | 0.00023 | 13.338 |
| LP Basal Health | No | 0.8852 | 0.0223 |

*Supporting Table 4: Summary statistics from the t-test and Fisher ratio analysis of the 11 metabolites of interest from four pathways.*

| Location | Status | Metabolite | Pathway | p-value | FDR p-value | Fisher Ratio |
|---|---|---|---|---|---|---|
| Apical | Healthy | valine | Amino Acid | 0.0128 | 0.1559 | 2.1265 |
| Apical | Healthy | serine | Amino Acid | 0.0257 | 0.2243 | 1.5781 |
| Apical | Healthy | leucine | Amino Acid | 0.0134 | 0.1605 | 2.0341 |
| Apical | Healthy | isoleucine | Amino Acid | 0.0143 | 0.1685 | 2.174 |
| Apical | Healthy | glycine | Amino Acid | 0.9979 | 0.9993 | 0 |
| Apical | Healthy | alanine | Amino Acid | 0.0688 | 0.3022 | 1.1978 |
| Apical | Healthy | lithocholic acid | Cholesterol | 0.0218 | 0.2168 | 1.4118 |
| Apical | Healthy | cholesterol | Cholesterol | 0.0031 | 0.0694 | 4.1786 |
| Apical | Healthy | glucose-6-phosphate | Glycolysis | 0.5659 | 0.7779 | 0.0626 |
| Apical | Healthy | glucose | Glycolysis | 0.2879 | 0.5728 | 0.2387 |
| Apical | Healthy | alpha-ketoglutarate | TCA Cycle | 0.5145 | 0.748 | 0.0991 |
| Apical | WD | valine | Amino Acid | 0.1613 | 0.3406 | 0.4384 |
| Apical | WD | serine | Amino Acid | 0.0292 | 0.2169 | 1.229 |
| Apical | WD | leucine | Amino Acid | 0.1128 | 0.2921 | 0.5917 |
| Apical | WD | isoleucine | Amino Acid | 0.0903 | 0.2737 | 0.6789 |
| Apical | WD | glycine | Amino Acid | 0.7069 | 0.7977 | 0.0281 |
| Apical | WD | alanine | Amino Acid | 0.612 | 0.7286 | 0.0501 |
| Apical | WD | lithocholic acid | Cholesterol | 0.7438 | 0.8173 | 0.0217 |
| Apical | WD | cholesterol | Cholesterol | 0.0002 | 0.0806 | 10.6946 |
| Apical | WD | glucose-6-phosphate | Glycolysis | 0.086 | 0.2695 | 0.6751 |
| Apical | WD | glucose | Glycolysis | 0.2865 | 0.4771 | 0.2285 |
| Apical | WD | alpha-ketoglutarate | TCA Cycle | 0.2779 | 0.4645 | 0.273 |
| Basal | Healthy | valine | Amino Acid | 0.8002 | 0.9851 | 0.0113 |
| Basal | Healthy | serine | Amino Acid | 0.9484 | 0.9965 | 0.0007 |
| Basal | Healthy | leucine | Amino Acid | 0.7702 | 0.9813 | 0.015 |
| Basal | Healthy | isoleucine | Amino Acid | 0.8187 | 0.9891 | 0.0092 |
| Basal | Healthy | glycine | Amino Acid | 0.2803 | 0.9125 | 0.2219 |
| Basal | Healthy | alanine | Amino Acid | 0.9472 | 0.9965 | 0.0008 |
| Basal | Healthy | lithocholic acid | Cholesterol | 0.4944 | 0.9341 | 0.0861 |
| Basal | Healthy | cholesterol | Cholesterol | 0.5394 | 0.9436 | 0.0706 |
| Basal | Healthy | glucose-6-phosphate | Glycolysis | 0.6495 | 0.9555 | 0.0367 |
| Basal | Healthy | glucose | Glycolysis | 0.8161 | 0.9891 | 0.0098 |
| Basal | Healthy | alpha-ketoglutarate | TCA Cycle | 0.6433 | 0.9499 | 0.0382 |
| Basal | WD | valine | Amino Acid | 0.1574 | 0.8473 | 0.4058 |
| Basal | WD | serine | Amino Acid | 0.0981 | 0.8473 | 0.5649 |
| Basal | WD | leucine | Amino Acid | 0.1914 | 0.8473 | 0.3677 |
| Basal | WD | isoleucine | Amino Acid | 0.093 | 0.8473 | 0.6356 |
| Basal | WD | glycine | Amino Acid | 0.6949 | 0.9367 | 0.0278 |
| Basal | WD | alanine | Amino Acid | 0.7393 | 0.9457 | 0.0196 |
| Basal | WD | lithocholic acid | Cholesterol | 0.3677 | 0.8473 | 0.1545 |
| Basal | WD | cholesterol | Cholesterol | 0.1784 | 0.8473 | 0.3612 |
| Basal | WD | glucose-6-phosphate | Glycolysis | 0.5045 | 0.9192 | 0.0806 |
| Basal | WD | glucose | Glycolysis | 0.6359 | 0.9367 | 0.0397 |
| Basal | WD | alpha-ketoglutarate | TCA Cycle | 0.9106 | 0.9875 | 0.0022 |

### Supporting Figures

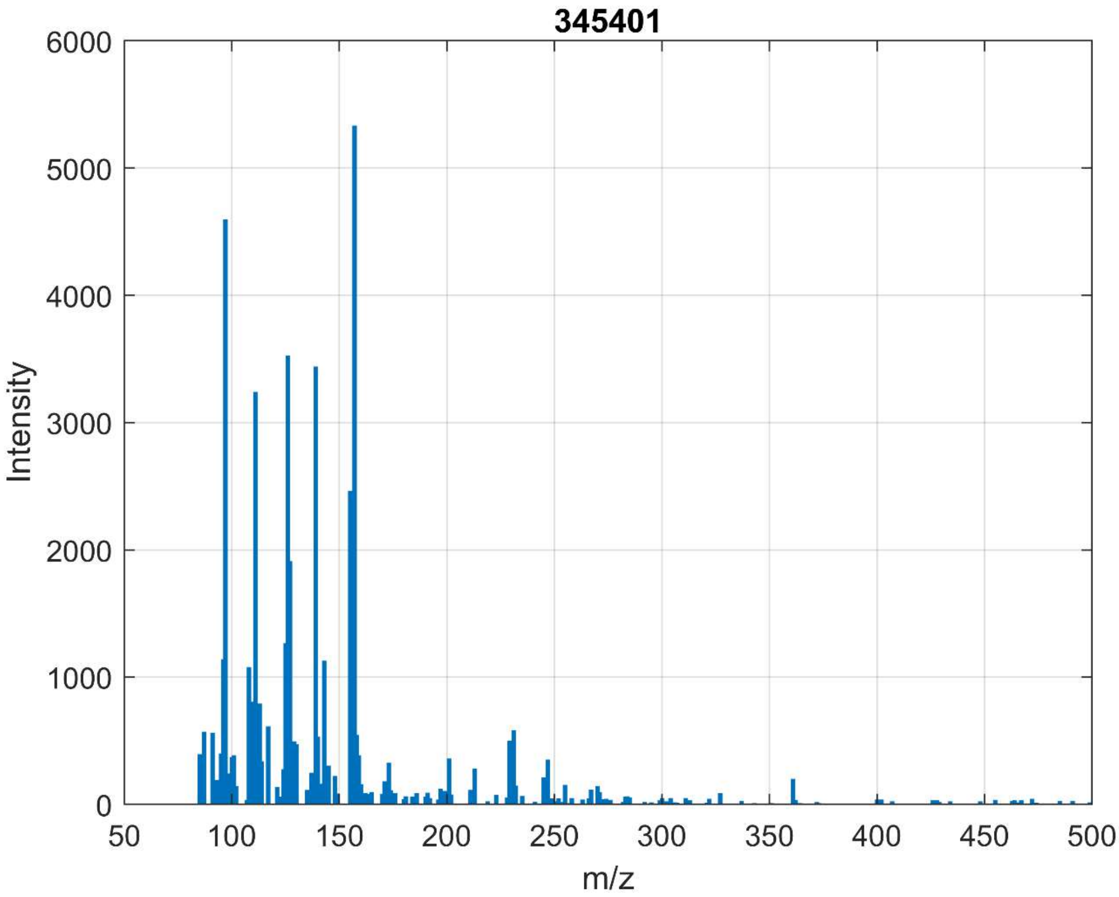


*Supporting Figure 1: Mass spectrum of metabolite 345401, a chlorine containing metabolite identified as overrepresented in dapagliflozin treated samples. Its structure remains unknown.*

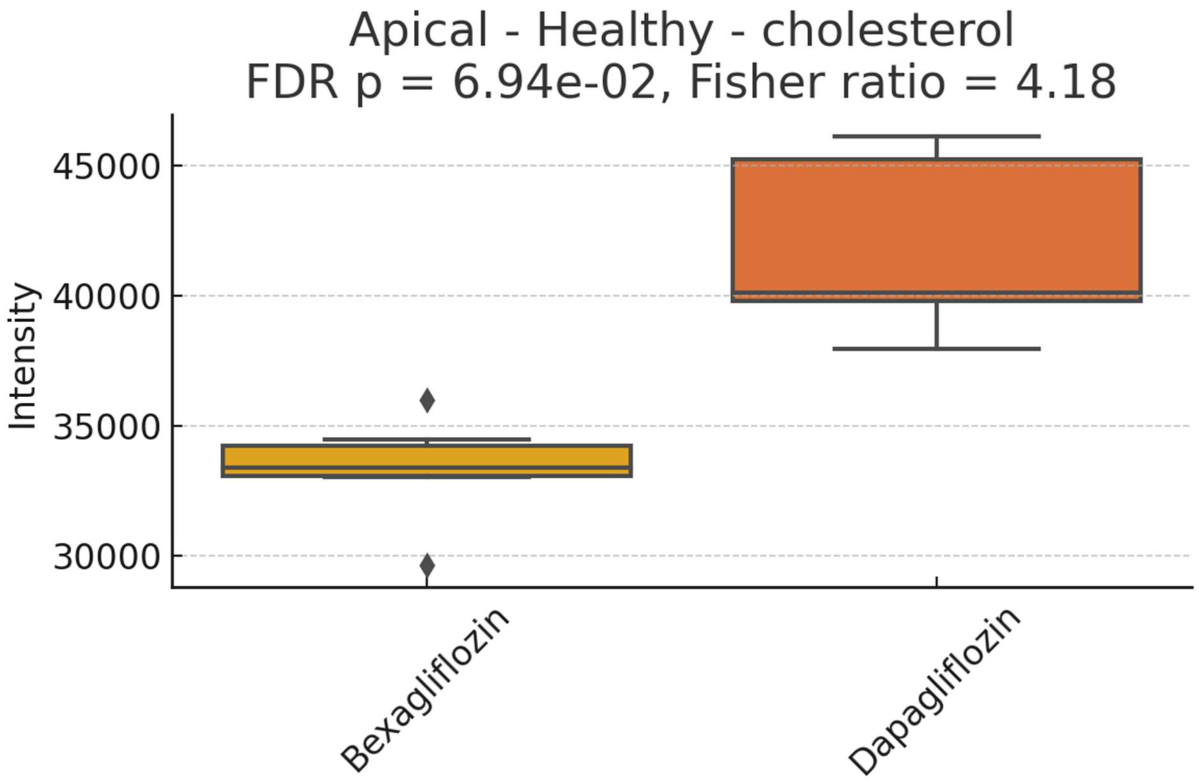


*Supporting Figure 2: Box plot of relative abundances of cholesterol in apical, healthy cells, treated with either bexagliflozin or dapagliflozin.*

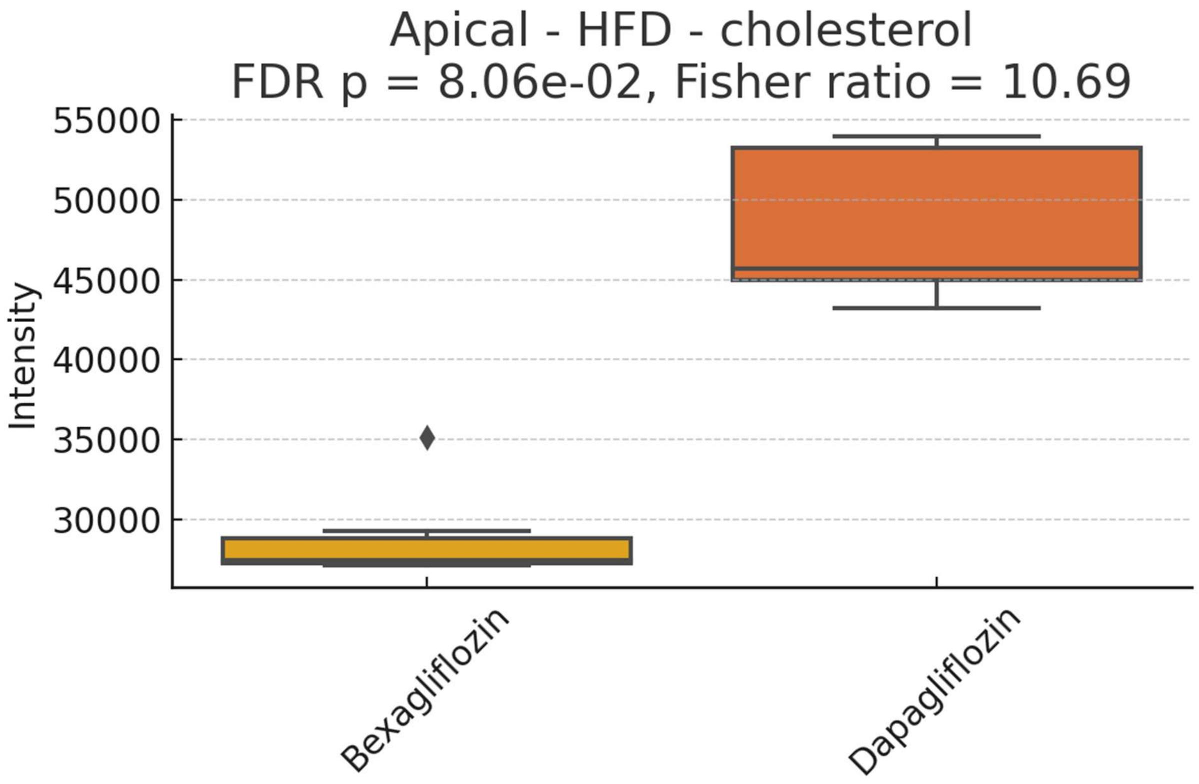


*Supporting Figure 3: Box plot of relative abundances of cholesterol in apical, WD cells, treated with either bexagliflozin or dapagliflozin.*

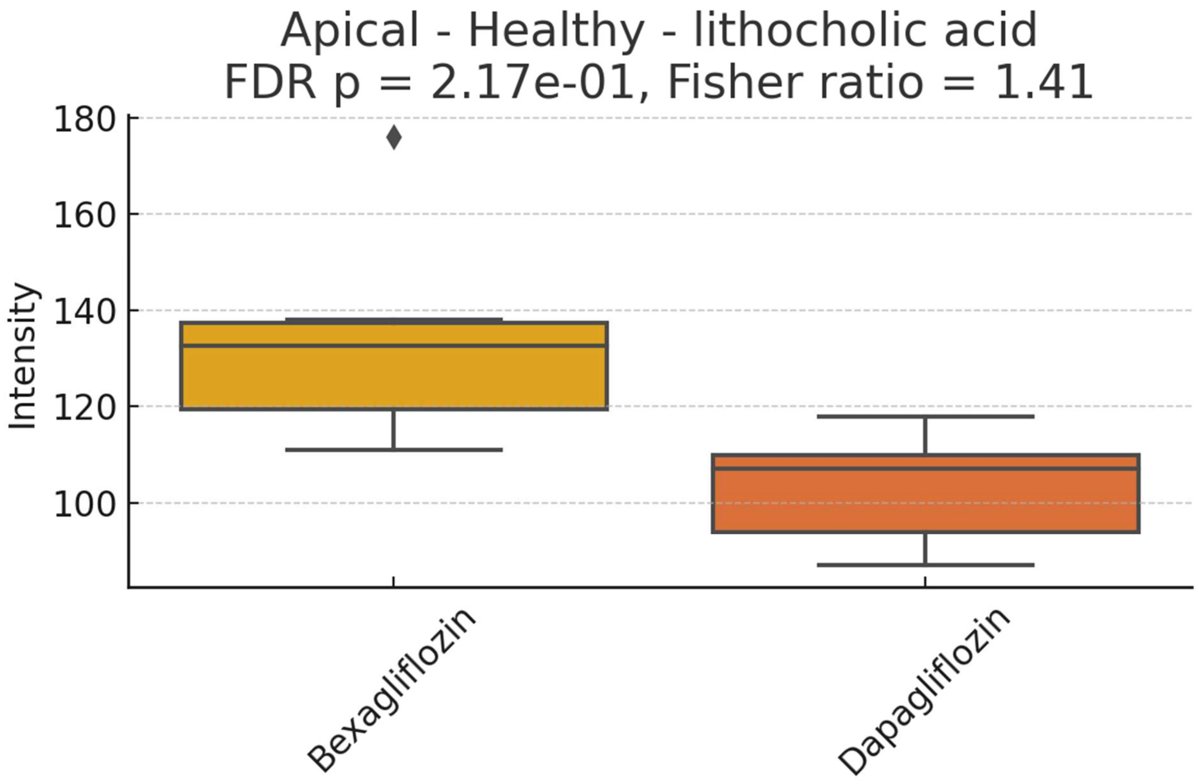


*Supporting Figure 4: Box plot of relative abundances of lithocholic acid in apical, healthy cells, treated with either bexagliflozin or dapagliflozin.*
